# Extinction risk predictions for the world’s flowering plants to support their conservation

**DOI:** 10.1101/2023.08.29.555324

**Authors:** Steven P. Bachman, Matilda J.M. Brown, Tarciso C.C. Leão, Eimear Nic Lughadha, Barnaby E. Walker

## Abstract

- More than 70% of all vascular plants lack conservation status assessments. We aimed to address this shortfall in knowledge of species extinction risk by using the World Checklist of Vascular Plants to generate the first comprehensive set of predictions for a large clade: angiosperms (flowering plants, ∼330,000 species).
- We used Bayesian Additive Regression Trees (BART) to predict the extinction risk of all angiosperms using predictors relating to range size, human footprint, climate, and evolutionary history and applied a novel approach to estimate uncertainty of individual species level predictions.
- From our model predictions we estimate 45.1% of angiosperm species are potentially threatened with a lower bound of 44.5% and upper bound of 45.7%.
- Our species-level predictions, with associated uncertainty estimates, do not replace full Red List assessments, but can be used to prioritise predicted threatened species for full Red List assessment and fast-track predicted non-threatened species for Least Concern assessments. Our predictions and uncertainty estimates can also guide fieldwork, inform systematic conservation planning and support global plant conservation efforts and targets.

## Introduction

There is an urgent need to ‘bend the curve’ of biodiversity loss (Mace *et al*., 2018), as indicators report widespread and ongoing decline (Tittensor *et al*., 2014; Haddad *et al*., 2015; Díaz *et al*., 2019). Ambitious goals and targets have been set to stem this loss, achieve stabilisation, and build a path to recovery, whilst ensuring human needs are fairly met (Milner-Gulland *et al*., 2021; Gumbs *et al*., 2021; CBD, 2021b). Progress against these goals and associated targets will be measured using a range of robust biodiversity indicators (CBD, 2021a,b,c). However, clades such as plants present considerable challenges for assessment and monitoring due to their ubiquity (Bar-On *et al*., 2018), diversity (Brummitt *et al*., 2021), data gaps and bias (Meyer *et al*., 2016).

The conservation status of species, often expressed as their risk of extinction, is an example of a fundamentally important metric that is challenging to assess and monitor at the scale of major clades such as plants, ∼350,000 vascular plant species. Despite long being a policy target of the plant conservation community (see Global Strategy for Plant Conservation – Target 2)(Sharrock *et al*., 2018), a comprehensive conservation status assessment for all plants remains an outstanding knowledge gap (Bachman *et al*., 2018; Nic Lughadha *et al*., 2020). Shortfalls in coverage of conservation assessments for plants are most pronounced when considering the gold standard for assessments – the IUCN Red List of Threatened Species™ (hereafter Red List)(IUCN, 2012). Of the known ∼350,000 vascular plant species, the latest version of the Red List (IUCN, 2022) documents 62,666 species (∼18%), a sample that is subject to taxonomic, geographic and trait biases (Nic Lughadha *et al*., 2020). One approach to address inherent human bias in deciding which species to assess is to select a random sample for assessment, as used for the Sampled Red List Index (Brummitt *et al*., 2015). Although useful for estimating the overall proportion of species that are threatened, major threat drivers, and trends over time (Tittensor *et al*., 2014), the sampled assessments only provide a small contribution (1,000s of species) to overall coverage. The remaining 82% of species that do not have a Red List assessment are important because the Red List has become the *de facto* resource for extinction risk assessments. Red List assessments have been mainstreamed into numerous conservation schemes such as Key Biodiversity Areas (IUCN, 2016), Species Threat Abatement and Restoration (Mair *et al*., 2021), and Tropical Important Plant Areas (Darbyshire *et al*., 2017). Ambitious goals for area-based conservation such as the 30/30 challenge (Dinerstein *et al*., 2019; Maxwell *et al*., 2020) must be supported with comprehensive coverage of extinction risk assessments to ensure that conservation potential is maximised.

Recent methodological advances (Walker *et al*., 2021), targeted funding (Botanic Gardens Conservation International, 2021), and availability of custom tools (Bachman *et al*., 2011; Brummitt *et al*., 2015; Moat, 2020) are all helping to address the plant Red List assessment shortfall. A recent review (Cazalis *et al*., 2022) noted two broadly defined approaches currently being employed to fast-track assessments: i) *criteria-specific* and ii) *category-predictive*. Criteria-specific approaches use automated procedures to calculate specific parameters such as extent of occurrence (EOO) that enable application of Red List criteria, while in category-predictive approaches Red List categorisations such as ‘Endangered’ or ‘Threatened’ are predicted using correlative modelling techniques, often using the current Red List as a model training set. Category-predictive approaches have been employed in several recent analyses and show promise for rapid estimation of Red List categories for plants (Darrah *et al*., 2017; Nic Lughadha *et al*., 2019; Zizka *et al*., 2021; Walker *et al*., 2021; Silva *et al*., 2022), but have yet to be applied to achieve comprehensive coverage at the scale of a large clade such as plants. Furthermore, these approaches are being carefully scrutinised to ensure consistency (Walker *et al*., 2020) and to recommend best practice (Walker *et al*., 2021).

With this growing body of evidence that machine learning and modelling techniques can be useful for predicting extinction risk in large clades of unassessed species, it is timely to consider a comprehensive study at the scale of a group like plants. Vascular plants (∼350,000) are predominantly made up of angiosperms, the flowering plants, with 335,798 species (Govaerts *et al*., 2021; Brown *et al*., 2023b). The Gymnosperms (conifers, cycads and relatives) have been near-comprehensively assessed for the Red List and are well supported by IUCN Red List specialist groups for assessment updates, hence there is no demand for extinction risk predictions. The limited and skewed samples of Ferns and allies published on the Red List (6% assessed) makes it hard to train models with sufficient accuracy to be useful. We therefore focus this study on generating predictions for angiosperms.

There are limiting factors in any attempt to predict extinction risk such as the quality of the training set used to build models. The IUCN Red List represents the ‘gold standard’ dataset for knowledge on extinction risk (Rodrigues *et al*., 2006; Betts *et al*., 2020; Cazalis *et al*., 2022) and has several characteristics that make it suitable for modelling angiosperms: balance of class size (42% threatened vs 58% non-threatened); size (62,666 species); representativeness of group being predicted (see Supplementary Figure S1) and accuracy of dataset. The latter is supported through provision of detailed guidelines to assess species for the Red List, quality checks, expert review, and a petitions process that all contribute to a robust and consistent dataset (IUCN Standards and Petitions Committee, 2022).

Accurate prediction of extinction risk requires identifying and obtaining an appropriate set of predictor variables. Previous studies, covering different taxonomic groups, have reported high predictive performance when using predictors of extinction risk such as restricted geographic range size, exposure to human threats, evolutionary history, presence in particular habitats or biomes and life history traits (Bland *et al*., 2015; Silva *et al*., 2022; Caetano *et al*., 2022). When determining appropriate predictors, there is often a trade-off between resolution (predictors supported by high-resolution data) and completeness (predictors with data coverage for all/most species), especially when dealing with species-rich groups such as plants. For example, two high-resolution predictors of geographic distribution that are used directly in Red List assessments such as extent of occurrence (EOO) and area of occupancy (AOO) have proven to be useful for predicting extinction risk (Pelletier *et al*., 2018; Zizka *et al*., 2021; Bellot *et al*., 2022). However, measuring EOO and AOO is not possible for most plant species due to inherent bias and gaps in the occurrence data used to calculate them (Meyer *et al*., 2016). Use of these high-resolution predictors is therefore often at the cost of excluding a large proportion of data-poor species, thereby introducing several systematic biases that are non-independent from extinction risk (Meyer et al 2016). However, comprehensive and inclusive coverage of geographic distribution at a coarser scale is now possible for all vascular plants with the publication of the World Checklist of Vascular Plants (Govaerts *et al*., 2021)(WCVP). The WCVP dataset includes species-level data such as life form classification and native distributions recorded at the resolution of level 3 in the World Geographical Scheme for Recording Plant Distributions (WGSRPD) (Brummitt et al., 2001), hereafter ‘botanical countries’. For instance, the number of ‘botanical countries’ in a species’ native range is known for all vascular plants, and, although coarser in resolution, is a good proxy for EOO (see Supplementary Figure S2). Predictors derived from the WCVP are harnessed in this study to achieve comprehensive extinction risk prediction coverage of angiosperm species.

Several machine learning approaches for predicting extinction risk have already been tested and compared across regions and taxa (Nic Lughadha *et al*., 2019; Walker *et al*., 2021), and both Random Forests and Neural Networks have consistently provided high levels of performance. However, whilst most studies explore the ability of models to generate accurate predictions on unseen data through evaluation statistics such as accuracy and true skill statistic (TSS), few studies have reported uncertainty at the species level for final classifications (although see Silva et al., 2022). Uncertainty estimates for extinction predictions are valuable when converting predictions into full, global, Red List assessments. Species predicted with high certainty of being threatened can be treated as high priorities for full Red List assessment and species predicted with high certainty of being non-threatened can be fast-tracked for Red List publication with automated tools such as Rapid Least Concern (Bachman *et al*., 2020). Clusters of ‘problematic’ species with highly uncertain predictions could be priorities for further fieldwork and ground-truthing thereby helping to reduce data deficiency and improve future iterations of comprehensive extinction risk predictions. Systematic conservation planning can also use these predictions to fill gaps where full Red List assessments are not available and can explicitly incorporate uncertainty (Tulloch *et al*., 2013). To provide informative predictions with uncertainty, we use a Bayesian Additive Regression Trees (BART) modelling framework that has performed well with classification tasks and allows for the quantification of uncertainty (Chipman et al., 2010). To our knowledge, this approach has not previously been used for extinction risk predictions.

Our aims are therefore to use BART to predict extinction risk for all angiosperm species and quantify the uncertainty of each species-level prediction. Using these predictions, we estimate the overall percentage of species threatened, including uncertainty, and guide the prioritisation of species for full assessments for the Red List. We explore the performance of these models, the importance of the predictor variables, and possible reasons for misclassifications to inform future iterations of these models. We provide a pipeline from which to reproduce these results and update predictions as part of the ongoing monitoring of global plant extinction risk.

## Materials and methods

### Data sources and processing

A list of angiosperm species was obtained from the World Checklist of Vascular Plants (WCVP)(Govaerts *et al*., 2021; Brown *et al*., 2023b). We filtered the WCVP using the list of families from the Angiosperm Phylogeny Group IV (2016) to remove non-angiosperms from our species list. We selected all species recognised as accepted by WCVP and removed hybrids to leave a dataset with 328,565 species. Species’ native distributions are recorded at the resolution of ‘botanical countries’, (Brummitt *et al*., 2001), and were filtered to exclude species occurrences coded as introduced, extinct or doubtfully present.

The latest version of the Red List IUCN Red List of Threatened Species (2022.2) was downloaded from the Red List website (https://www.iucnredlist.org/) in July 2022 using the search terms: Taxonomy = Plantae - Kingdom, Geographical Scope = Global, which included 62,666 species. From these species, 60,231 were matched to the WCVP using exact and fuzzy matching functions from the *rWCVP* package (Brown *et al*., 2023b). The species categorised as Extinct (EX, 112 species), or Extinct in the wild (EW, 40 species) were removed and the obsolete ‘lower risk’ categories were mapped to their relevant categories in version 3.1 of the IUCN Red List Categories and Criteria (IUCN, 2012). The Data Deficient (DD) species were grouped with the Not Evaluated (NE) species to give 275,004 unassessed species. The remaining 53,512 data sufficient (CR, EN, VU, NT, LC) species were used to train our model.

### Predictor calculation

Predictors were selected to represent five important correlates of extinction risk: geographic distribution (number of botanical countries), evolutionary history (first 50 eigenvectors capturing evolutionary relatedness among species), traits (lifeform), human threats (human footprint values from 2009 (10 classes) and change in human footprint values from 1993 to 2009 (5 classes) (Venter *et al*., 2016)), and biomes (16 biomes from the WWF Ecoregions dataset (Olson *et al*., 2001), including rock & ice and lakes). In addition, we included the year of description, derived from WCVP, as a predictor based on the finding that recently described plant species are more likely to be threatened (Brown *et al*., 2023a). See Table S3 for the full list of predictors and item S4 for a description of the predictors and how they were generated. The number of botanical countries was log transformed, species without lifeform information (24%) were coded as ‘unknown’ and all other predictors with missing values were imputed with a k-nearest neighbour function using Gower distances (Gower, 1971).

### Model training and evaluation

We trained a Bayesian Additive Regression Trees (BART) model, to predict the extinction risk of all unassessed angiosperm species. A BART model is based on an ensemble of decision trees, and fits the sum of outputs to the data (Chipman *et al*., 2010). The BART model also limits overfitting by placing regularising priors on parameters that determine how individual trees are grown, constraining them to fit a small subsection of the data (weak learners). During fitting, values for these parameters are sampled using a Markov Chain Monte Carlo (MCMC) algorithm, resulting in a Bayesian posterior distribution of trees that can be sampled to provide uncertainty estimates for all predictions.

There were 53,512 assessed angiosperm species (non-data deficient) and 275,004 un-assessed angiosperm species (Data Deficient or Not Evaluated). Species categorised as Least Concern (LC) or Near Threatened were grouped as ‘non-threatened’ and species categorised as Vulnerable (VU), Endangered (EN), or Critically Endangered (CR) were grouped as ‘threatened’. Applying IUCN reporting standards (IUCN Red List Committee, 2022) gives an estimated proportion threatened of 42%, with a lower bound of 39% and upper bound of 47% when considering best- and worst-case scenarios for Data Deficient species.

We used nested cross-validation to both tune the hyper-parameters of our models and evaluate the resulting performance, splitting the species into five outer folds with three inner folds each. We generated the outer folds of this nested cross-validation scheme in two ways: by randomly excluding a fifth of species from each training set (random cross-validation); by randomly excluding entire families from each training set so that roughly a fifth of the assessed species were held out in each fold (family-wise block cross-validation). This splitting scheme allowed us to tune the BART hyperparameters on each set of inner folds, select the best performing hyperparameters for each outer fold, and evaluate the performance of a model trained using these hyperparameters on the corresponding test set of each outer fold. We chose these schemes to evaluate how well the model predicts the extinction risk of species similar to those in the training data and the extinction risk of species in families that have not yet been assessed for the IUCN Red List, respectively.

We trained a BART model on the training set of each CV fold and selected a classification threshold that maximised the ROC-AUC (Receiver Operating Characteristics - Area Under the Curve). We then evaluated these models on the corresponding test set, assessing their performance using TSS. The TSS is a performance metric that balances the proportion of threatened species correctly predicted as such (sensitivity) with the proportion of non-threatened species correctly predicted (specificity). The TSS ranges from 1 for perfect predictions to -1 if there are no correct predictions. For a model that predicts all species as non-threatened, or all species as threatened, the TSS will be 0 (sensitivity of 1 + specificity of 0 - 1 = 0). We split the performance scores across different subsets of data to see how well extinction risk was predicted for features such as lifeforms or biomes. We carried out exploratory data analysis on any misclassified species to identify potential explanations for model misclassifications (such as certain threats not being adequately captured in our predictors).

After evaluating the performance of the models, we trained our final model on all species with (non-data deficient) Red List assessments, using the classification threshold that gave the highest ROC-AUC on this training set, to account for the higher proportion of not threatened species in the training dataset.

To explore potential bias associated with assessments on the Red List that are considered out of date (> 10 years since assessment), we trained a separate model using only assessments from the last ten years and recorded performance metrics.

Finally, we explored how well the model will perform against future assessments. Using the same methodology as described above, we trained a BART model on the previous version of the IUCN Red List (version 2022-1) and evaluated its predictions for the newly published assessments in version 2022-2.

### Predicting extinction risk

We used this final model to predict the extinction risk (threatened or not threatened) of all 275,004 unassessed angiosperm species. Predictions can be made from a BART model either by drawing from the posterior predictive distribution, giving direct predictions of threat, or by drawing from the posterior probability that a species is threatened. We used each approach for a different purpose. We used 1,000 samples from the posterior predictive distribution for each species to generate aggregate predictions of the proportion of threatened species across all angiosperms and different sub-groups of angiosperm, quantifying uncertainty as the width of the 95% credible interval. We used 1,000 samples from each species’ posterior probability of being threatened to classify that species as threatened if the mean posterior probability was above our classification threshold, flagging predictions where our threshold was within the 95% credible interval as low confidence (uncertain).

We explored patterns of extinction risk in angiosperms by visualising predicted extinction risk against various features derived from the WCVP dataset such as taxonomy, lifeform and biome. We mapped the spatial distribution of angiosperm extinction risk to explore the impact of the predictions on patterns of threatened species compared to the existing Red List. We filtered the final predictions based on those species that were currently unassessed for the global Red List, predicted to be threatened and above our threshold for certainty. These potential priority species were mapped to highlight areas where plant Red List assessment activity is urgently needed. Finally, we looked at how our model predicted threat status for species currently listed as Data Deficient on the Red List.

## Results

### Model classification performance

The overall accuracy of our model was within the bounds of those reported in previous studies adopting a variety of modelling techniques (Zizka *et al*., 2021; Walker *et al*., 2021; Silva *et al*., 2022; Bellot *et al*., 2022) (Table 1 and Table S7). Average sensitivity was 0.76 and average specificity was 0.81. Sensitivity was marginally lower than specificity, indicating that when the model was wrong, the error was more likely to be an underprediction of a threatened species rather than an overprediction of a non-threatened species. Average TSS was 0.58 and average accuracy was 0.79. The exclusion of older assessments (>10 years) had minimal effect on model performance and model performance on unseen assessments (2022-2 assessments not included when trained on an earlier model which was trained on version 2022-1) was similar to our final model (Table S8).

**Table 1.**
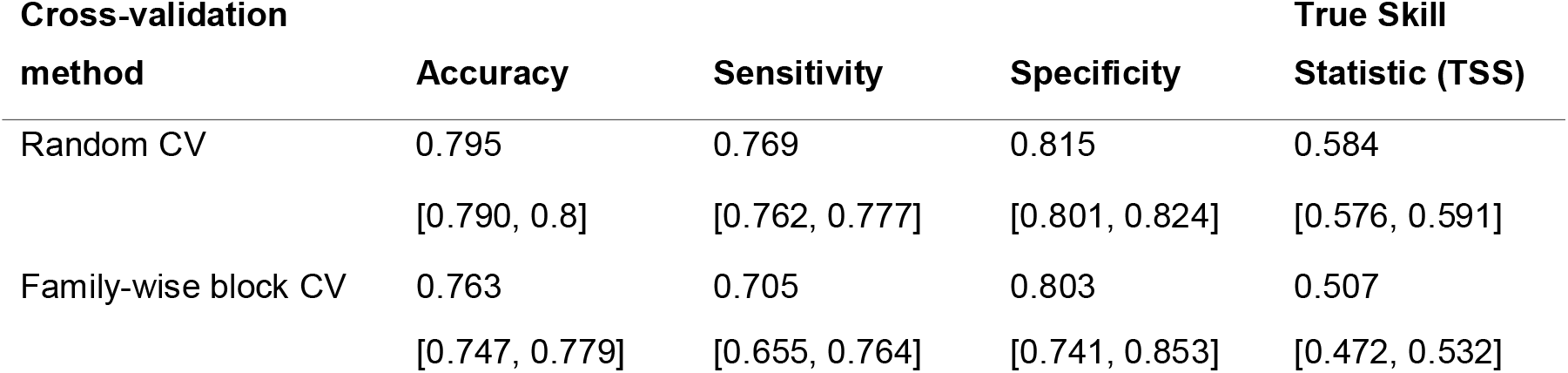
Performance statistics based on random and family-wise block cross-validation. Average values for each statistic, followed with values for 95% credible intervals from the 5-fold cross-validation in square brackets.

Certain groups within the dataset were better predicted than others, for example the threat status for annuals was more accurately predicted than that of other lifeforms, and species from tropical biomes generally had more accurate predictions than temperate or sub-tropical biomes (Figure S9). The random cross-validation performance scores were marginally but consistently higher than those of family-wise block cross validation.

The TSS values for species occurring in botanical countries across Africa, southern America, and tropical Asia were moderate to good (average 0.39 – 0.49). Parts of Europe had high TSS scores e.g., Austria, Hungary, Germany, and Great Britain (>0.80). In contrast, some regions such as Northern America and Temperate Asia (except Kazakhstan, 0.66 TSS) had low average TSS values (0.12 – 0.27 respectively; Figure S10A). Both Northern America and Temperate Asia had good specificity scores, indicating that the low sensitivity scores (Table S11) i.e., the reduced ability to predict threatened species, were pulling down the overall TSS. Average sensitivity decreased as species geographic distribution increased, showing an expected difficulty in predicting large-ranged species as threatened (Figure S12).

We further explored the reasons for low sensitivity (misclassifying threatened species as non-threatened) by examining the specific IUCN Red List criteria cited in the assessments of threatened species that were misclassified as non-threatened. Noting that at least one criterion must be cited for each species assigned to a threatened category and many assessments cite more than one criterion, Criterion B (small geographic range) is the most frequently cited criterion overall (76%), followed by D (small populations, 16%), A (population reductions, 12%) and C (small and declining populations, 5%) (Figure S13). Notably, the sensitivity is lower when criterion A is cited compared to when it is not cited. Criterion B includes two metrics to describe distributions, the area of occupancy (AOO) and the extent of occurrence (EOO). We explored whether the inclusion of either or both metrics impacted the sensitivity scores (Figure S14). Sensitivity was slightly higher for species citing criterion B1 (EOO) than those citing criterion B2 (AOO), but the highest sensitivity was achieved when both were cited. Species with relatively large ranges (i.e., larger than the threshold for Criterion B) were more likely to be misclassified, highlighting the importance of range size in our model.

Finally, we looked at threat coding for assessed threatened species to see if there were particular threats for species that our models misclassified. If there was a particular threat type e.g., invasive species, with many more species that were incorrectly classified compared to correctly classified, this could indicate that we are missing this threat process from our models. For most threats, the proportion of species classified correctly and incorrectly was similar, suggesting our models are not systematically excluding an important threatening process affecting plants (Figure S15). The following five threats were marginally overrepresented in the misclassified threatened species: Residential & commercial development, Transportation & service corridors, Natural system modification, Invasive & other problematic species, and Pollution. Conversely, Biological resource use, which was not included in the scope of our predictors, was cited for a higher proportion of correctly classified species.

### Angiosperm extinction risk

Based on our extinction risk predictions (Table S16), the percentage of angiosperm species threatened was 45.1% (Figure 1, with 95% credible intervals from 44.5% to 45.7%), which was marginally higher than the mid-point estimate calculated from the latest Red List (42%, 39 – 47% lower and upper bounds (IUCN Red List Committee, 2022)). We explored the proportion of predicted threatened species in different sub-groups across all angiosperms. We found that epiphytes are the most threatened lifeform and annuals are the least threatened. Threat percentage of epiphytes increased from 49% based on assessed species from the Red List to 53.9% with predictions (95% CI between 52% and 56%).

**Figure 1.**
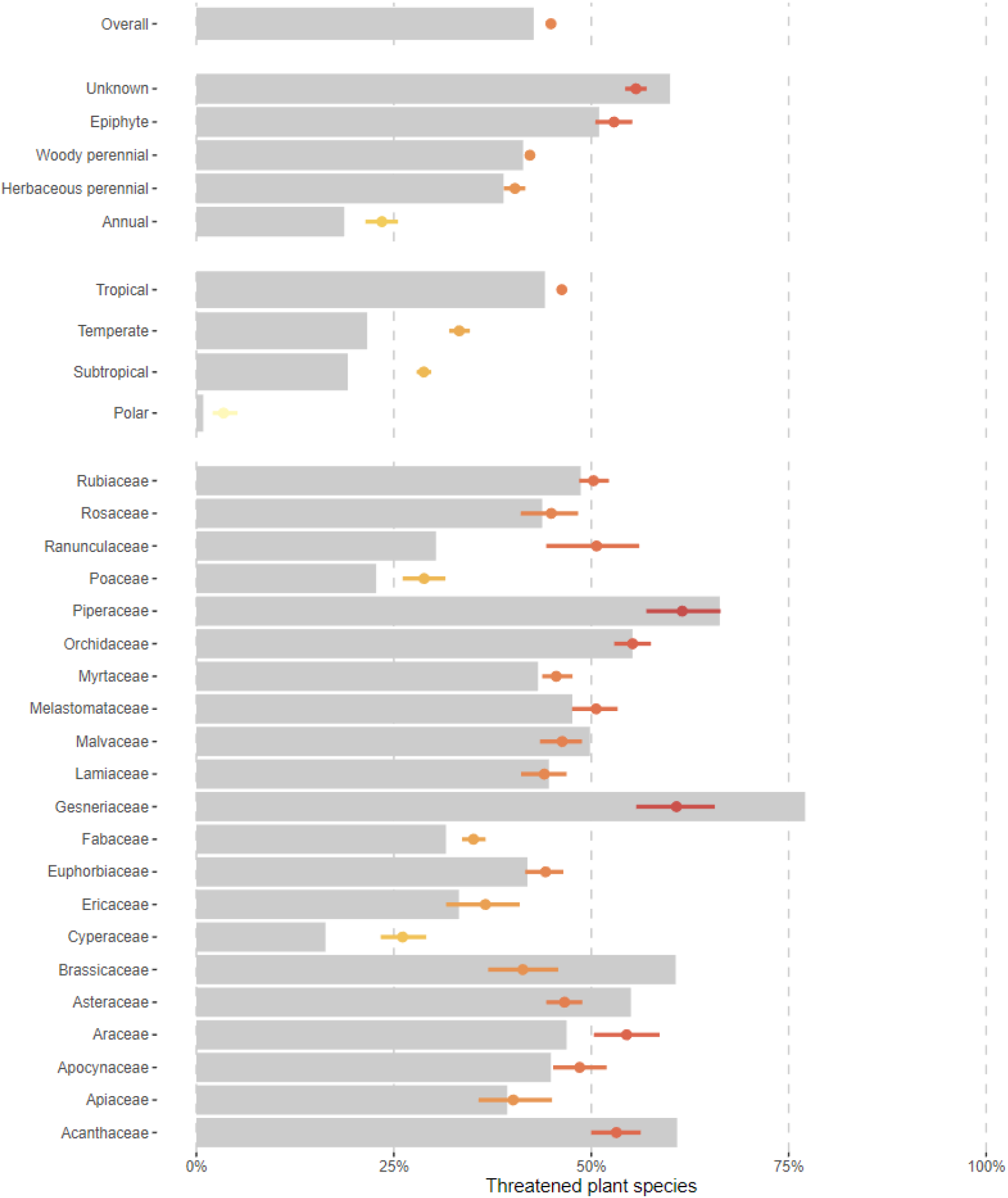
Proportion of species predicted as threatened (coloured points; lines show 95% credible interval) in comparison to proportion of species assessed as threatened on the Red List (grey bars) for different facets of the dataset. All species (A), lifeforms (B), biomes and largest families (C, filtered on families with at least 3,000 species). Where credible intervals are small (<2%) the coloured lines are not visible.

Among the 24 largest plant families (>= 3,000 species), the predicted five most threatened families were Piperaceae (60% threatened), Gesneriaceae (58%), Bromeliaceae (56%), Orchidaceae (56%), Araceae (55%). The predicted proportion of threatened species per family was generally in line with observed values from Red List, but some families were exceptions e.g., Araceae, Cyperaceae, Ericaceae, Poaceae and Ranunculaceae were predicted to be more threatened than on the Red List. In contrast, Acanthaceae, Brassicaceae, Caryophyllaceae, and Gesneriaceae were predicted to be less threatened than observed from the Red List. Gesneriaceae was noticeable in being one of the most threatened families from predictions, but not as threatened as observed from the current Red List. Predictions of biomes (tropical, subtropical, temperate and polar) were in line with the Red List with the tropics being the most threatened. Of the 4,505 species categorised on the Red List as Data Deficient (DD), our model predicted that 71% (3,212 species) are threatened, of which 77% were predictions with high certainty. Individual species predictions with uncertainties are provided in Table S16.

Geographic spread of predicted threat at the scale of botanical countries (Figure 2A) broadly reflects those observed from the current Red List (Figure 2C and 2E). The areas of highest predicted threat were broadly congruent with observed data, with islands and archipelagos such as Hawaii, Madagascar, New Caledonia, Borneo, and the Philippines amongst the highest percentage threatened. Uncertainty was highest amongst smaller islands, Iceland, Sweden, and Finland, but otherwise was evenly spread across botanical countries (Figure 2B). There are large gaps in Red List assessment coverage across western parts of North America, southern parts of South America, Europe, Temperate and South-East Asia, and Australia (Figure 2D). Central and northern South America, South-East Asia, Madagascar and South Africa had the highest numbers of potential priority species (i.e. those that are unassessed and confidently predicted as threatened; Figure 2F).

**Figure 2.**
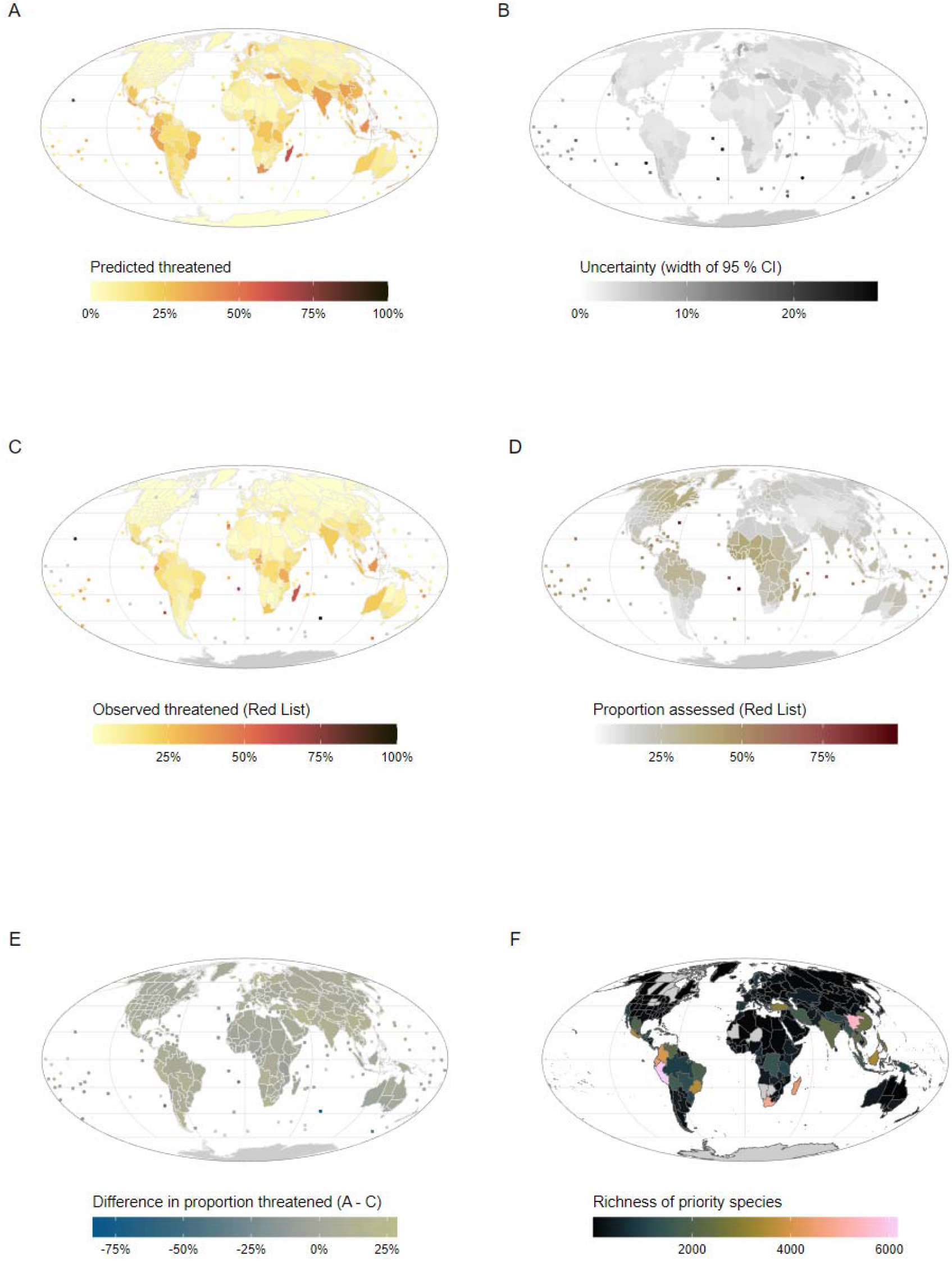
(A) The proportion of angiosperm species predicted to be threatened in each botanical country. (B) the corresponding uncertainty measured as width of the 95% credible interval on the posterior predictive for the proportion of threatened species in each botanical country. (C) Observed proportion of angiosperm species in each botanical country from the Red List version 2022-2. (D) Observed coverage of species assessed on the Red List as proportion of total number in each botanical country. (E) Difference between the predicted and observed proportion threatened (A – C). (F) Richness of potential priority angiosperm species based on all species currently unassessed for the Red List and predicted to be threatened with confidence.

### Importance of predictors

Of the 85 predictors included in the models, geographic distribution (represented here by number of botanical countries) was by far the most important single predictor (Figure S17). However, when grouped by the six main types of predictors (Figure 3), the importance of human footprint, year of description, biome and phylogenetic relatedness became more apparent, although geographic distribution measured in number of botanical countries was still the most important predictor. Geographic distribution, human footprint, biome, and phylogenetic relatedness all showed variation over the cross-validation folds. Lifeform was the least important factor contributing to model accuracy.

**Figure 3.**
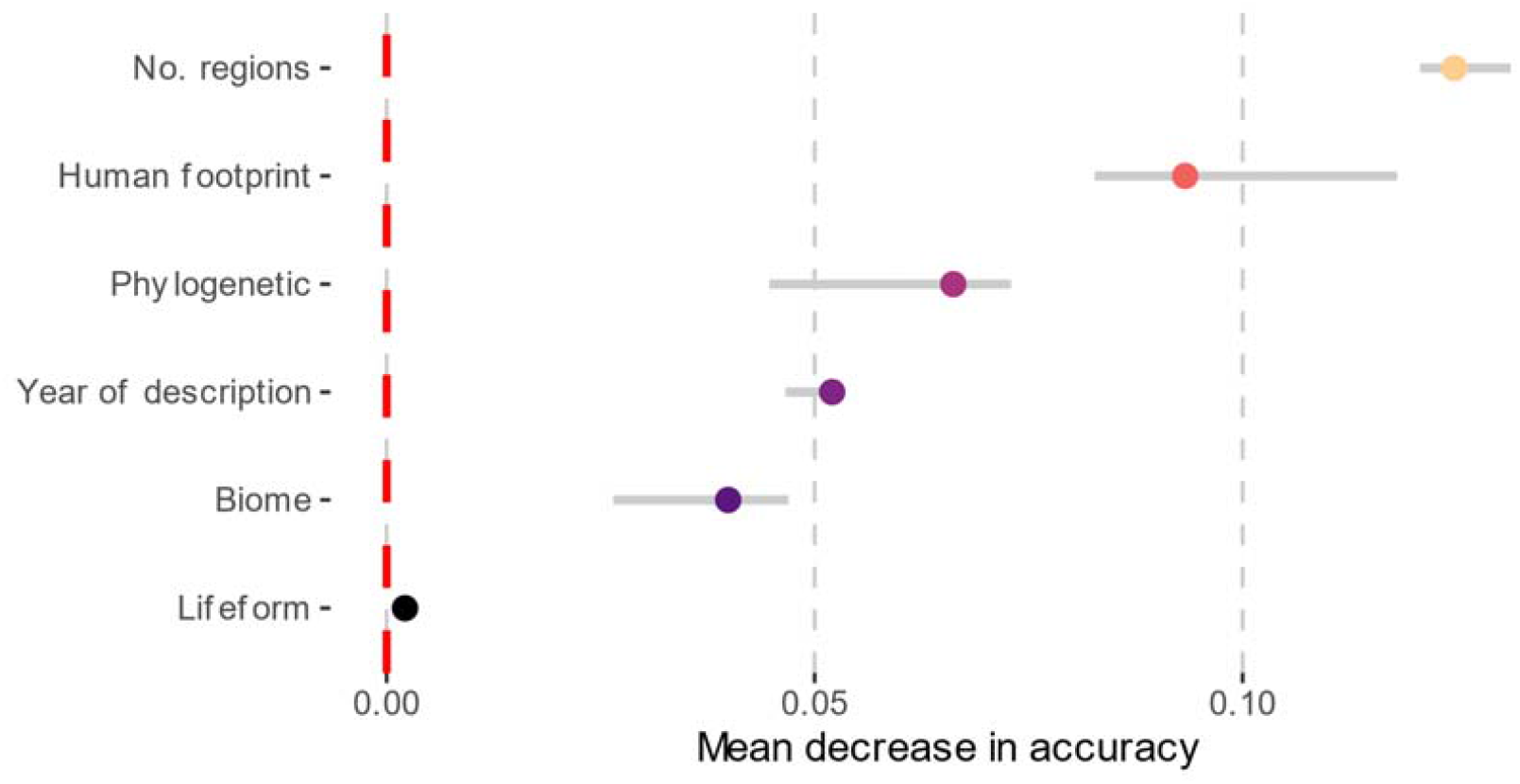
Predictor importance grouped by type of predictor, based on mean decrease in accuracy. No. regions = number of botanical countries. Phylogenetic = phylogenetic eigenvectors capturing evolutionary relatedness among species. Grey lines show 95% credible interval.

### Uncertainty in predictions

The classification threshold was 0.433, meaning species at or below this value were classified as threatened and species above this value were classified as non-threatened. Considering all species predictions, 15% (50,743) were classified as low certainty because the 95% credible intervals from the posterior distribution crossed the classification threshold for threatened. Conversely, 85% (277,773) were classified with certainty. See table S16 for individual species prediction probabilities and uncertainty values.

## Discussion

Our machine learning approach demonstrates we can generate comprehensive species-level estimates of extinction risk (and uncertainty) using coarse-scale predictors. We predicted extinction risk for all 328,565 known angiosperm species, with 79% accuracy validated on the latest published global Red List. Based on predictions only, we estimate 45.1% of angiosperms are threatened (95% CI 44.5% – 45.7%). This rapid and repeatable estimation of extinction risk provides an overview of plant extinction risk when formal assessments are unavailable, closing a substantial knowledge gap and providing a basis from which full Red List assessments could be prioritised.

### Model performance

Based on cross-validation, our models performed well, but patterns in both over- and under-prediction errors may have consequences if they are used to prioritise formal Red List assessments. The models displayed higher specificity than sensitivity, so fewer non-threatened species were misclassified as threatened (than vice versa). The performance was similar for random versus family-wise cross-validation, but family-wise was more variable. Some families are not represented by species assessments on the Red List and are therefore absent from the training set e.g. Rapataceae [96 species], Rafflesiaceae [48 species]. The model is likely to generalise well to these unseen families, but for some it will not perform as well. Performance is likely to improve as more species from unassessed families are added to the Red List. In terms of spatial heterogeneity of performance, there was notably low performance for parts of northern America and temperate Asia (except Kazakhstan), driven by low sensitivity in these areas i.e., the model underpredicted the extinction risk to threatened species. We found that misclassified species were mostly wide-ranged threatened species (Figure S11); in general, these species were assessed as threatened under Criterion A (population decline), rather than Criterion B (range size), suggesting that our model performs better at detecting species likely to trigger Criterion B than A. This is supported by the importance of range size (measured as number of botanical countries; Figure 3) in our model, consistent with previous studies (Darrah *et al*., 2017; Nic Lughadha *et al*., 2019; Walker *et al*., 2021; Bellot *et al*., 2022). For north America, potential practical impacts of underpredicting threatened species are reduced by the availability of comprehensive assessments from NatureServe. A recent review of USA trees showed NatureServe and IUCN Red List assessments were similar (Carrero *et al*., 2022). Our assessment predictions are therefore likely to be most useful in regions where there is a greater proportion of unassessed and potentially threatened species predicted with a higher level of certainty, such as central and northern South America and South-East Asia.

### Angiosperm extinction risk patterns

Our predictions estimated that 45.1% (CI 44.5% to 45.7%) of all angiosperms are threatened. We found support for known spatial hotspots of extinction risk such as Madagascar and Hawaii. We found that epiphytes are highly threatened, with an elevated extinction risk of 53.9% (95% CI 52 – 56%, compared to the current Red List 49%), which is consistent with findings that 45% of epiphytes have a high proportion of species with ‘vulnerably small ranges’ compared to other life forms (Leão *et al*., 2023). For large families (>= 3,000 species) the estimations of percentage threatened are broadly similar for the Red List and our predictions (Figure 1). Differences may be due to numbers of Data Deficient species; the family Araceae is predicted to be more threatened than estimated from the Red List where 35% of assessments are Data Deficient. In contrast, the family Gesneriaceae, which performed well in the model validation (Figure S9) is predicted to be less threatened than estimated from the Red List and has a smaller proportion Data Deficient (11%), but had higher than average proportion of low-confidence predictions (22%). Biomes had similar proportions of threatened species from our predictions compared to the Red List. Our model predictions suggest that Data Deficient species are more likely to be threatened than not threatened, with 69% predicted threatened, of which 86% are with high certainty. This is consistent with recent estimates that more than half (56%) of 7,699 species categorised as Data Deficient on the IUCN Red List may be threatened (Borgelt *et al*., 2022).

### Towards more accurate predictions

Our model performed well overall, but inevitably there are trade-offs between accuracy and scale. An advantage of the World Checklist of Vascular Plants is its complete (or near-complete) coverage of species, distributions and associated data such as lifeform (Govaerts *et al*., 2021). However, the coarse resolution of botanical countries results in low precision in the circumscription of species’ ranges, making fine-scale range size metrics such as area of occupancy (AOO) and extent of occurrence (EOO) impossible to estimate from WCVP. Although coarse-scale data from this same source have successfully been used to predict extinction risk of bulbous monocots (Darrah *et al*., 2017), these authors and others have demonstrated the value of using finer scale occurrence data to calculate AOO and EOO (Zizka *et al*., 2021; Walker *et al*., 2021; Silva *et al*., 2022; Bellot *et al*., 2022). However, at least three unique localities are required (after data cleaning) to calculate EOO, so many species lack sufficient occurrence data for these approaches. Imposing minimum data thresholds when predicting extinction can drastically reduce species coverage; half of all orchid species (Zizka *et al*., 2021), and more than half of land plants (Pelletier *et al*., 2018) had insufficient occurrence data to calculate predictors, leading to omission from analyses of many of the range-restricted species most likely to be threatened.

Year of description (Brown *et al*., 2023a) has not previously been used in plant extinction risk predictions, but made a useful contribution to our model (Figure 3). The relationship between year of description and probability of being threatened is related to range size, as plant species currently being described by taxonomists tend to be restricted to a single botanical country, 92% in 2020, compared to <50% in 1900 (Brown *et al*., 2023a). However, range size alone does not completely explain the predictive power of year of description, suggesting it may be capturing additional predictive power linked to rarity, decline, fragmentation or other Red List criteria, and it is this additional predictive power that is being leveraged in our model.

The extent to which our estimates of angiosperm extinction risk represent actual levels of extinction risk depends a large part on the ability of the training set, the IUCN Red List, to accurately reflect real-world extinction risk. Bias in the Red List, such as the tendency to apply criterion ‘B’ more frequently (Figure S18) may mean that true extinction risk is being over- or under-estimated. In biodiversity hotspots undergoing habitat loss, species may be eligible to qualify for criterion A (or C, D, E), but have not been assessed under these criteria due to lack of data on population size, or lack of essential parameters such as generation length, although numbers of assessments applying criterion A have increased (Figure S19). The potential bias of using a single criterion has yet to be fully quantified, but when different criteria combinations were examined, the use of Criterion A on its own was marginally less threatened than when Criterion B was used on its own (Figure S20). However, it should be noted that all five Red List criteria (A-E) were designed to detect the symptoms of endangerment and only one criterion needs to be met to categorise a species as threatened (Mace *et al*., 2008). Advances in mapping species’ habitat requirements (Lumbierres *et al*., 2022) and availability of datasets with high temporal resolution (annual updates) offer options to plant Red List assessors beyond criterion B, although there is still a dependency on life-history parameters such as generation length that may hinder wider application of criterion A.

Finally, our predictors did not fully capture the extent of documented threatening processes (Nic Lughadha *et al*., 2020) such as over-harvesting, invasive species, impact of disease or climate change/severe weather. Climate change, broadly perceived to be a significant threat, has yet to be coded as such for many published Red List assessments for plants (Nic Lughadha *et al*., 2020) (a notable exception being (Davis *et al*., 2012)). Characterising the potential impact of future climatic conditions on plant species is a non-trivial task, requiring sufficient data to consider multiple climate scenarios and compliance with specific Red List guidelines (IUCN Standards and Petitions Committee, 2022).

### Implications for plant conservation and the Red List

The threat predictions from this study are not equivalent to published IUCN Red List assessments and are not intended to replace them. However, as highlighted in other studies (Zizka *et al*., 2021; Silva *et al*., 2022; Bellot *et al*., 2022), predictions can inform conservation research and prioritisation when used carefully, e.g. when combined with phylogenies to generate preliminary Evolutionarily Distinct and Globally Endangered (EDGE) scores (Forest *et al*., 2018). Indeed, recent work has demonstrated that by excluding unassessed or poorly-known species, we may be underestimating the extent to which threatened species are a key element of global biogeographic patterns (Brown et al., in review). We envisage that our comprehensive extinction predictions can be used in a wide range of future studies, though we stress that it is crucial that the strengths, biases, uncertainties, and limitations of predictions, as discussed here and elsewhere (Walker *et al*., 2020, 2021; Bellot *et al*., 2022), are taken into account when using predictions for downstream analyses.

To close the gap on the unassessed angiosperm species for the Red List we envisage three scenarios where our predictions may be useful for prioritisation, especially in the absence of any other evidence-based threat assessment. First, Red List specialist groups and authorities, national Red List programs and taxonomic or thematic Red List assessment projects could prioritise their assessment efforts on species that our model predicts with high certainty to be threatened. This will ensure our efforts are directed to the species likely to be in greatest need of full Red List assessment. Secondly, for the species we predict likely to be non-threatened, we encourage the fast-tracking of ‘Least Concern’ Red List assessments using automation with tools such as Rapid Least Concern (Bachman *et al*., 2020). Finally, those species with uncertain predictions might be triaged as Data Deficient and could be priorities for further fieldwork and ground-truthing.

When using the predictions generated in this study, the consequences of incorrect predictions must be carefully considered; each species incorrectly predicted to be threatened could result in sub-optimal allocation of the limited resources for assessments. Conversely, species predicted to be non-threatened should not be assumed to meet the criteria for Least Concern; such an assumption could result in assessments that understate extinction risk (i.e. are ‘non-precautionary’). Thus, caution is needed if using these results for conservation prioritisation, particularly for when high uncertainty or low accuracy has been demonstrated (e.g., temperate Asia).

Assessment coverage for plant species on the Red List is growing but remains low at 18%. If the concept of ‘assessment’ is defined more broadly as any evidence-based, digitally-available assessments, at any scale, plant species coverage exceeds 28% (Nic Lughadha *et al*., 2020). We therefore fall well short of Target 2 of the Global Strategy for Plant Conservation, an “assessment of the conservation status of all known plant species”(CBD, 2012). To ensure all threatened plant species are documented on the IUCN Red List, we need to prioritise the remaining unassessed species. The 328,565 predictions of extinction risk for all angiosperms generated in this study provide complete coverage at species level, with a quantification of uncertainty for each prediction. In the absence of any other evidence-based assessment these predictions give an overview of plant extinction risk and have multiple uses including prioritisation for full Red List assessment, identifications of problematic species i.e. high uncertainty, use in future research on threatened plant species, and informing systematic conservation planning and fieldwork campaigns.

## Acknowledgments

We thank all assessors, reviewers, facilitators and compilers that build the IUCN Red List of Threatened Species and the Red List Unit, Cambridge for maintaining this vital dataset. We thank the James Hutton Institute for access to a high-performance computer (HPC) cluster. We thank all contributors and compilers of the WCVP dataset.

## Author contributions

SPB, MJMB, TCCL, ENL and BEW planned and designed the research. SPB wrote the initial draft of the manuscript with input from MJMB, TCCL, ENL and BEW. MJMB, TCCL, BEW and SPB compiled and analysed the data. All authors interpreted the data and contributed to the final version of this paper.

## Data availability

The R code used for data processing, analysis and visualisation is available at: https://github.com/stevenpbachman/WCVP_threat_predictions

To be useful, our extinction risk predictions and their associated uncertainty estimates need to be open and easily accessible; we plan to integrate our results into the WCVP dataset and related toolkits (e.g. *rWCVP* (Brown *et al*., 2023b)), and provide regular updates.

