## Supplementary material for "Extinction risk predictions for the world’s flowering plants to support their conservation"

Index

### Figure S1 – representativeness of Red List

We explored whether the Red List was a suitable training set for predicting extinction risk of Angiosperms. We checked for representativeness across three dimensions of data that were linked to our chosen predictors: – taxonomy (families), geography (WGSRPD L3 regions) and traits (lifeform).

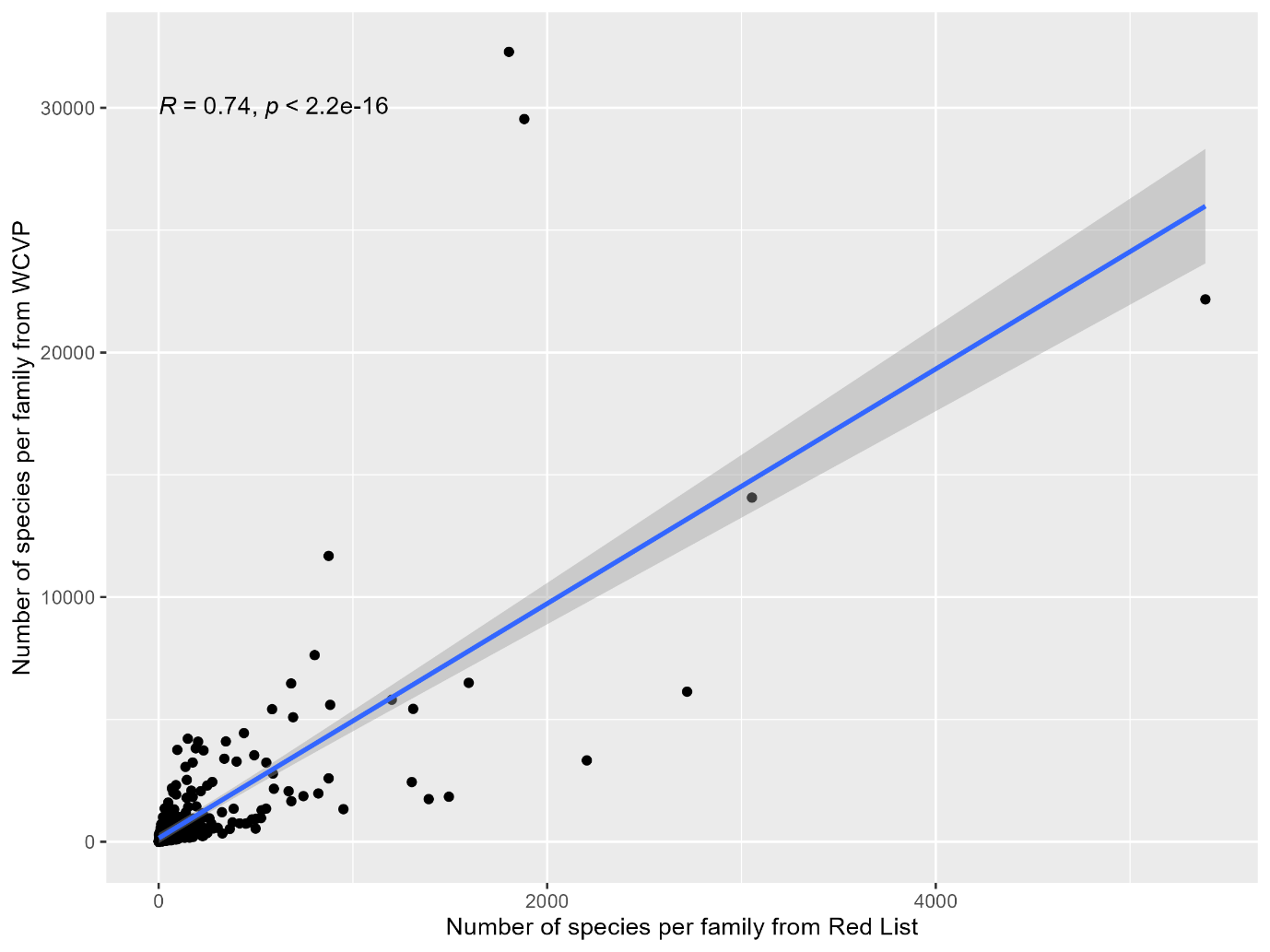

Figure S1A – correlation between number of species per family in the Red List vs number of species per family from WCVP Angiosperms

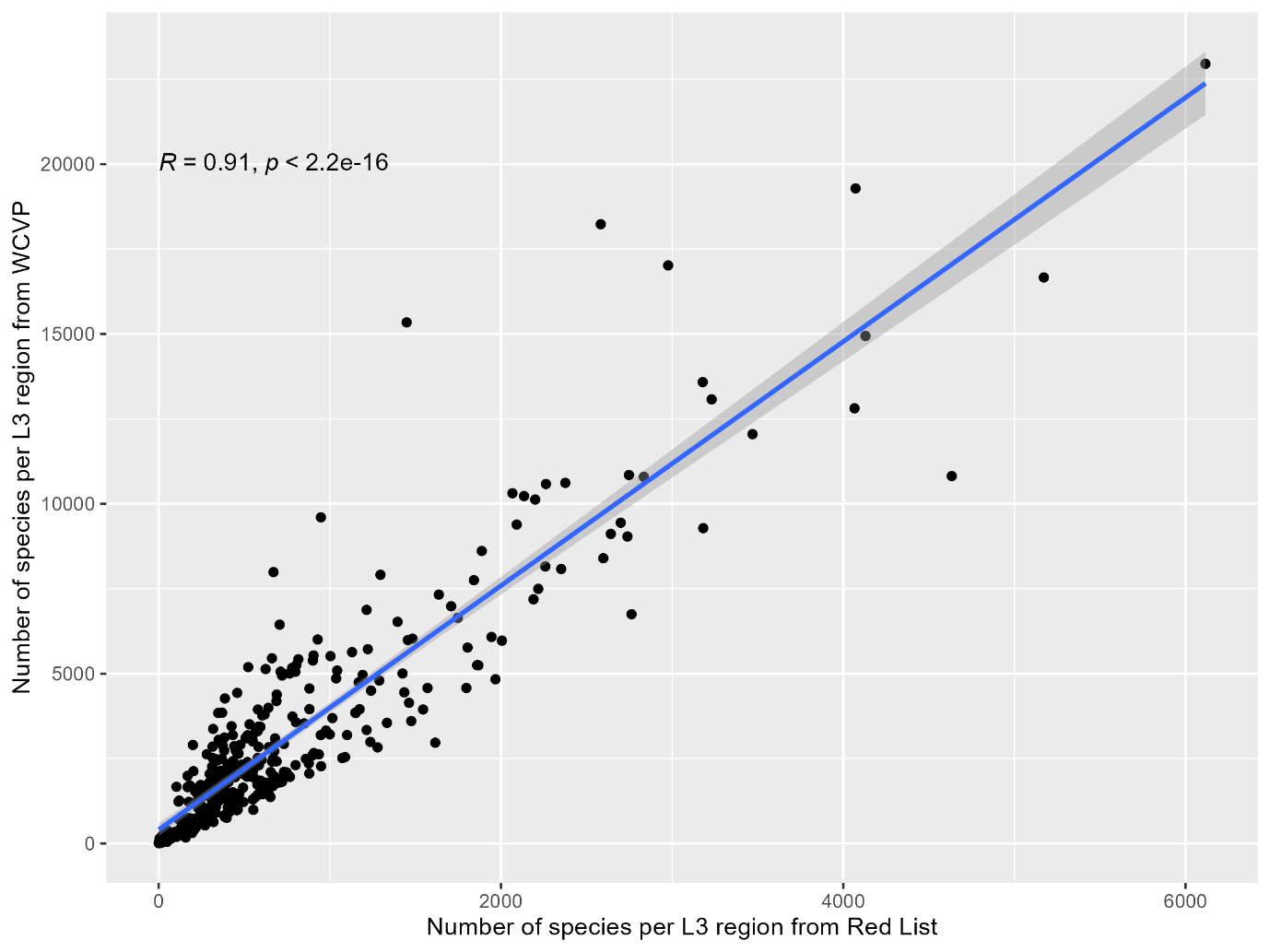

Figure S1B – correlation between number of species per L3 region in the Red List vs number of species per L3 region from WCVP Angiosperms

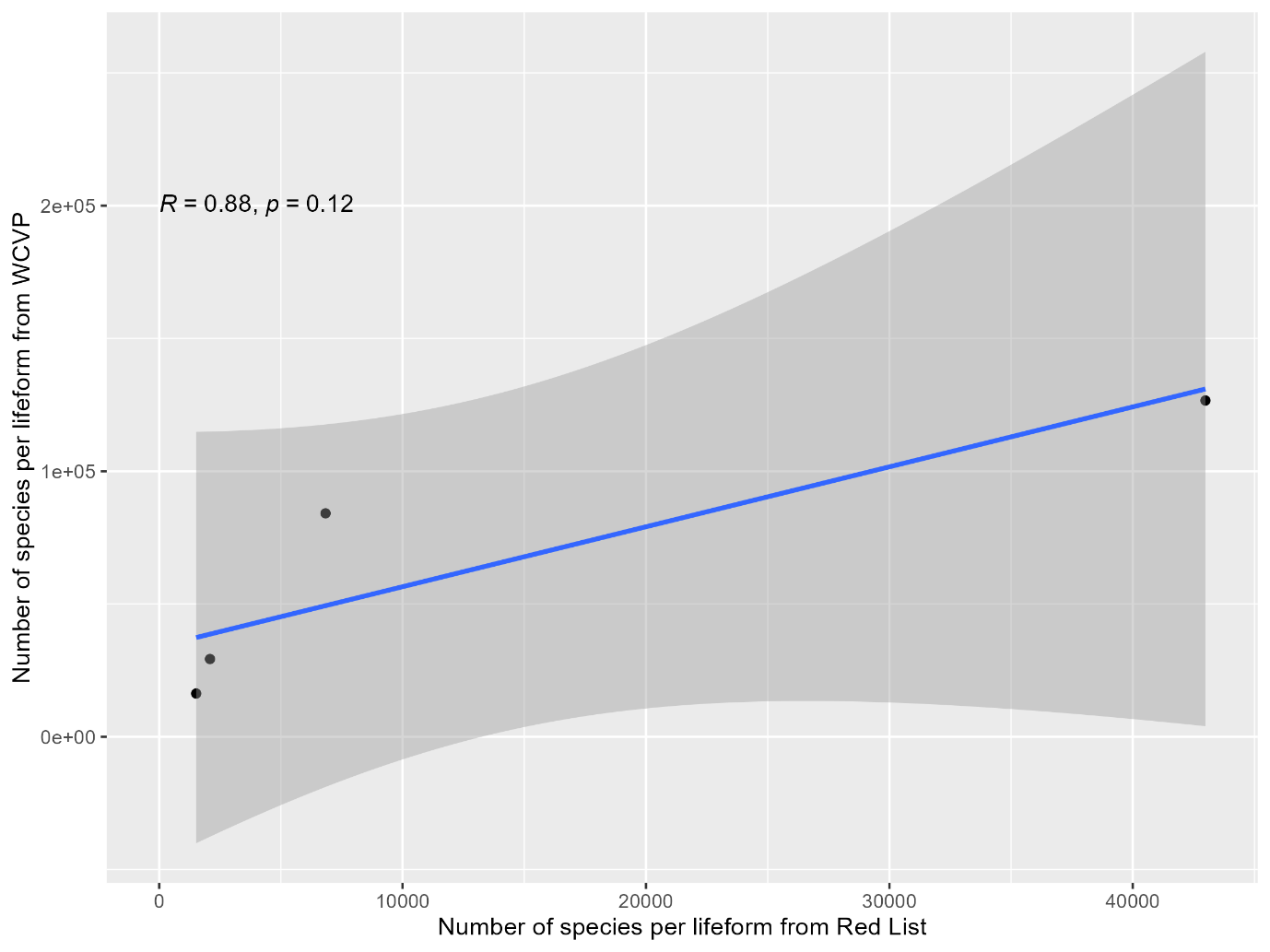

Figure S1C – correlation between number of species per Humphreys summarised lifeform class in the Red List vs number of species per Humphreys summarised lifeform class from WCVP Angiosperms

### Figure S2 – number of WGSRPD vs EOO

We used the number of WGSRPD regions as a proxy for the geographic range metrics used in the IUCN Red List criteria such as extent of occurrence (EOO) and area of occupancy (AOO) because EOO and AOO are not known for all Angiosperms. We compared number of WGSRPD regions (x-axis) vs extent of occurrence (EOO y-axis) and half of the variance in EOO was explained by the count of WGSRPD. Results for AOO were similar (not shown).

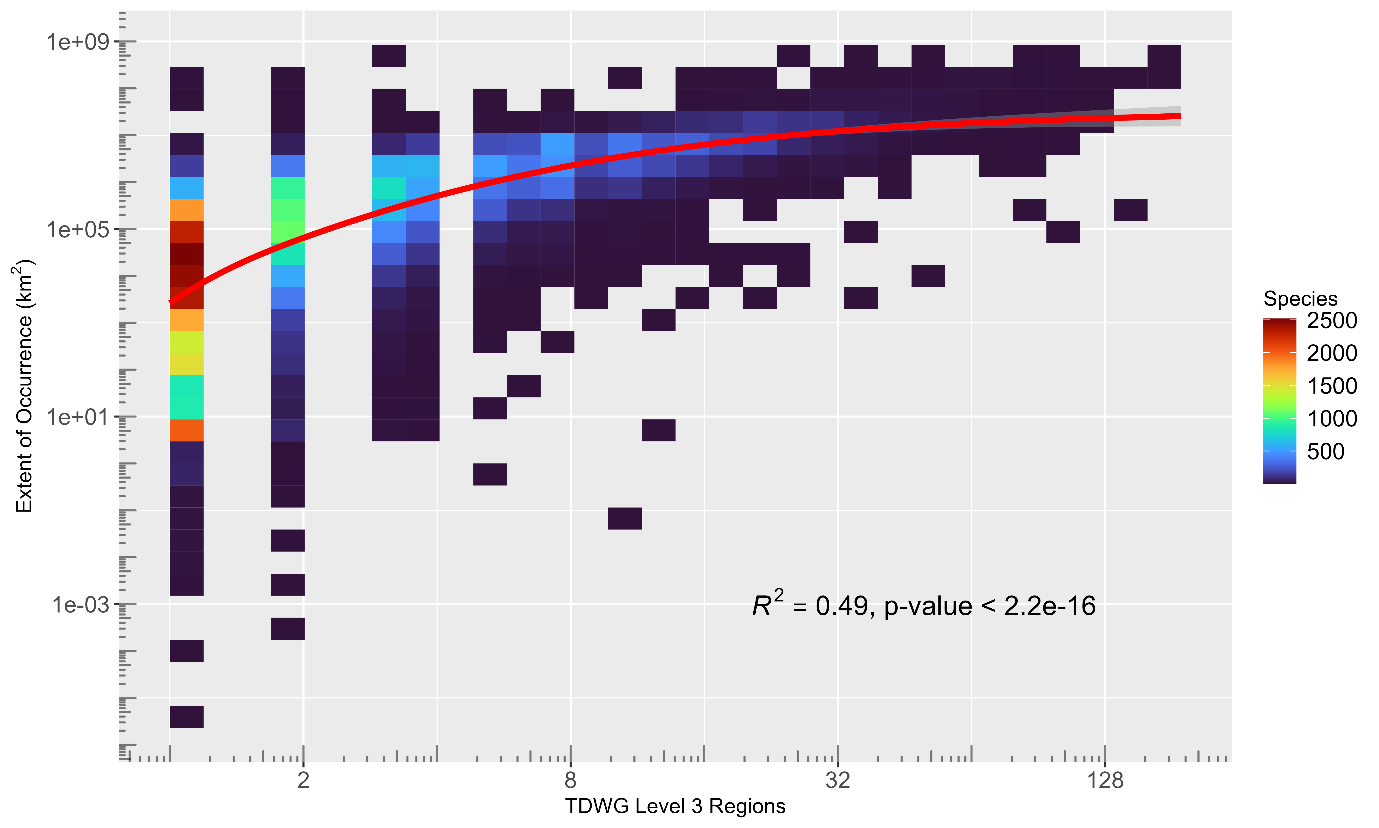

### Table S3 - list of predictors

See table **S3_list_of_predictors.csv** for the full list of predictors including category of predictor, description, data coverage, and source.

### Item S4 - description of predictor generation

The comprehensive coverage of angiosperm species in the World Checklist of Vascular Plants and associated geographic areas (World Geographic Scheme for Recording Plant Distributions) formed the basis of the model predictors.

#### Geography

Number of botanical countries was derived by summarising the number of unique native WGSRPD Level 3 regions each species occurred in.

#### Traits

Lifeform was taken from the WCVP dataset and was transformed to simplify and reduce the number of classes following. We adopted the same approach as used by (Humphreys et al., 2019) with modification by (Nic Lughadha et al., 2020– see Table S5 – life_form_mapping.csv.

#### Description Year

We used the first_published field from the WCVP dataset, which documents the year of publication of the taxon name, enclosed in parentheses. We removed parentheses and where species had a basionym we replaced the year with the year of publication of the basionym.

#### Phylogeny

Previous studies point to phylogenetic signals in the species’ geographic range sizes and extinction risk (Leão et al., 2014). Herein, we aimed to use the evolutionary similarities between closely related taxa to improve our predictions of extinction risk. We used the phylogenetic eigenvector method (Diniz-Filho et al., 2012, 1998), as it allowed us to incorporate the phylogenetic dependencies as a set of predictor variables in our machine learning models. Our first step was to obtain a comprehensive angiosperm phylogeny, up to date with the taxonomy of the World Checklist of Vascular Plants (Govaerts *et al*., 2021). We used the angiosperm phylogeny constructed by (Forest, 2023) Forest *et al*. (in prep) which updates the (Smith and Brown, 2018) GBMB tree according to recent changes and the taxonomy of the World Checklist of Vascular Plants. We simplified the tree to the genus level to reduce the dimensionality of the phylogenetic distance matrix to a manageable level. On this genus-level phylogeny, we extracted the pairwise phylogenetic distances between genera (i.e., phylogenetic distance matrix), which was transformed and double-centred before being used to extract the eigenvalues and eigenvectors representing the phylogenetic structure. The transformation and specific calculations were performed according to the ‘PVRdecomp’ function of the R package ‘PVR’ version 0.3(Santos, 2018). As the resulting number of eigenvectors was too large to be handled by the prediction models, we selected the eigenvectors containing the largest amount of phylogenetic information. We attempted a broken-stick method (Borcard *et al*., 2011), which selected 252 phylogenetic eigenvectors representing 77% of the variance associated with the phylogenetic tree. However, due to computational limitations, we used only the first 50 eigenvectors in the prediction model, representing 61.3% of the variance associated with the genus-level phylogeny tree (Figures S4.1).

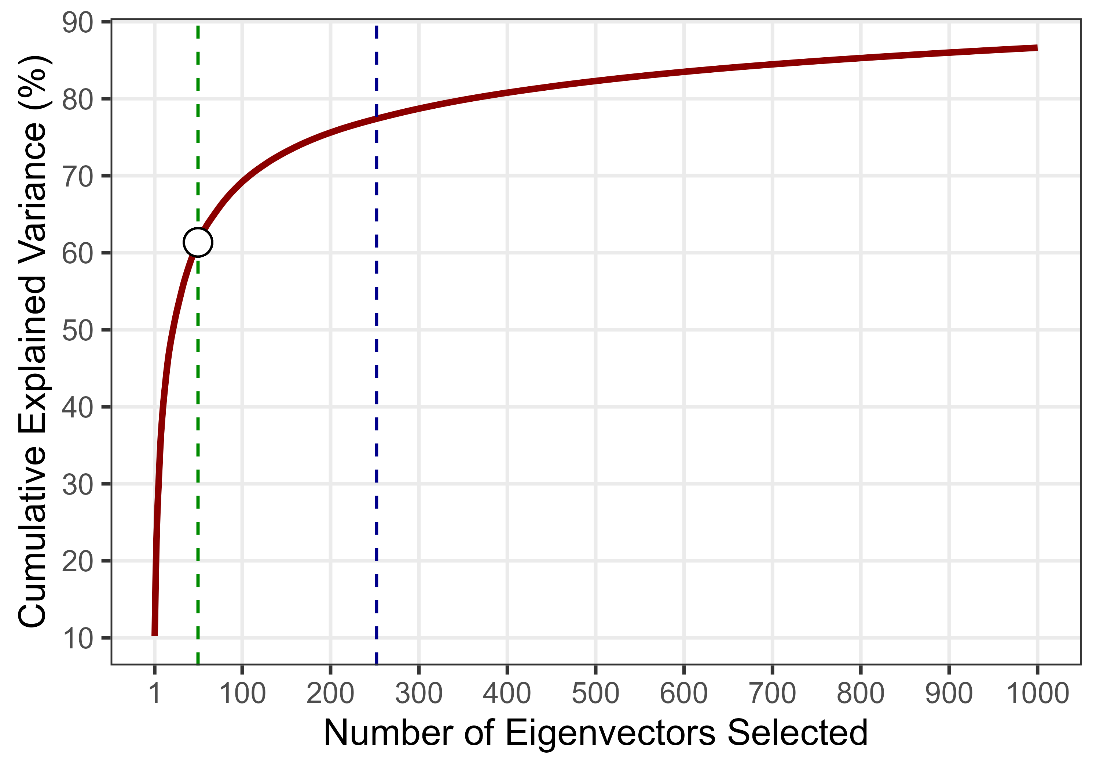

**Figure S4.1.** The cumulative variance of the genus-level angiosperm phylogeny captured by the first 1,000 phylogenetic eigenvectors. The vertical dashed blue line indicates the cutting point for the eigenvectors selected using the broken-stick method (i.e., the first 252 vectors, explaining 77% of the variance), while the green dashed line indicates the cutting point for the 50 eigenvectors (explaining 61% of the variance) we end up using due to computational limitation.

#### Biomes

We aggregated the WWF Ecoregions shapefile (Olson *et al*., 2001) to biome level (e.g. Tropical and Subtropical Moist Forests, Tundra). We calculated the area of each biome in each botanical country (Fig S4.2). Where the spatial polygons did not align perfectly (Fig S4.3a), we applied a correction factor so that the relative proportions of each biome remained the same, but the sum of these areas matched the total area of the botanical country (Fig S4.3b). Four botanical countries (all small, oceanic islands) did not appear in the ecoregion shapefiles; we manually estimated the biomes for these (Table S4.4).

***Figure S4.2.*** *Distribution of biomes across WGSRPD Level 3 Areas/botanical countries. WGSRPD Level 3 Areas/botanical countries are outlined; fill colour shows the biome from Olson et al. (2012).*

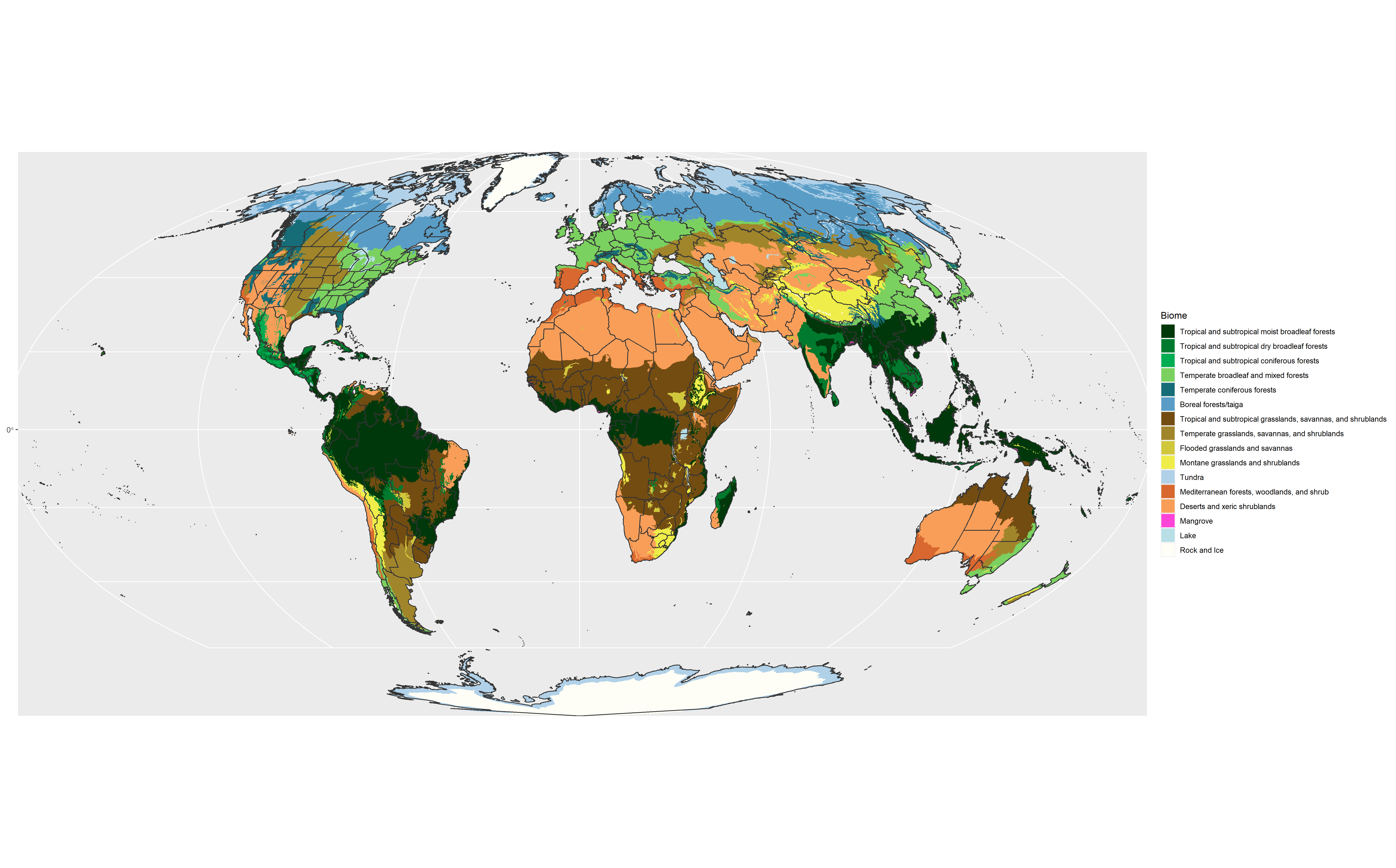

| **Table S4.4 Manual corrections for WGSRPD biomes** | | |
| --- | --- | --- |
| **CODE** | **Name** | **Biome** |
| MCI | Mozambique Channel Islands | Tropical and subtropical moist broadleaf forests (100%) |
| MCS | Marcus Island/Minami-tori Shima | Tropical and subtropical dry broadleaf forests (100%) |
| NRU | Nauru | Tropical and subtropical moist broadleaf forests (100%) |
| SEL | Selvagens Islands | Mediterranean forests, woodlands, and shrub (100%) |

***Figure S4.3.*** *Examples of discrepancies between ecoregions and WGSRPD shapefiles. A real example (the Falkland Islands) is shown in (a). Parts of the WGSRPD polygon that are not covered by ecoregions are highlighted in red, showing that the changes are due to different coastline resolutions. Resolution of this issue is shown in the fictional example (b): the relative proportions of Biomes 1 and 2 are retained, but the raw areas (in km^2^) are corrected so that the sum of the biome areas matches the total area.*

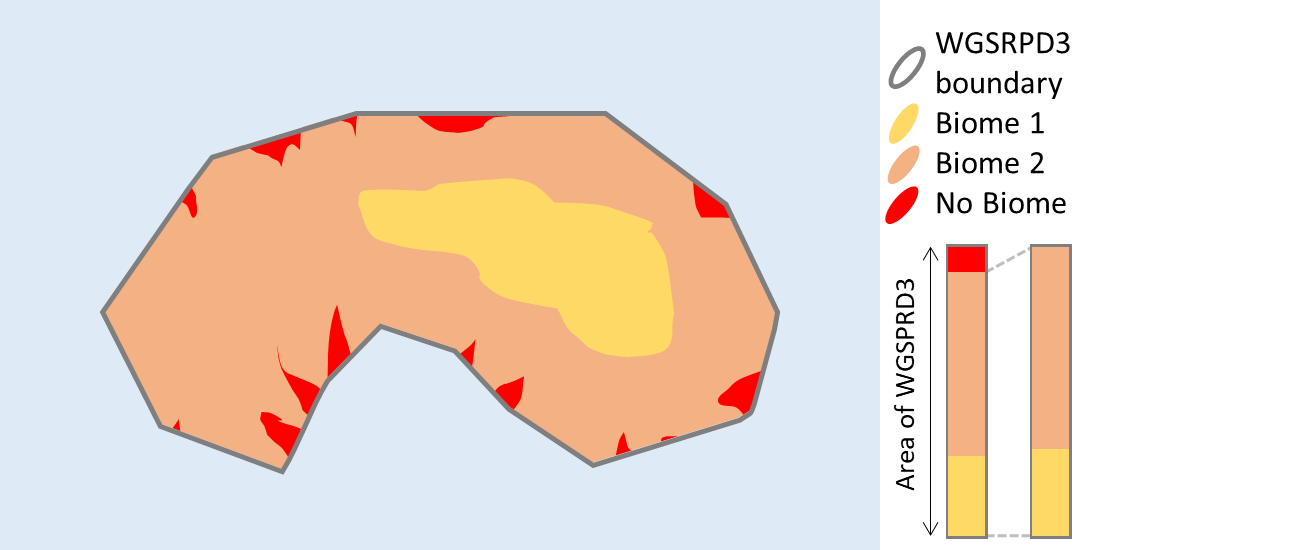

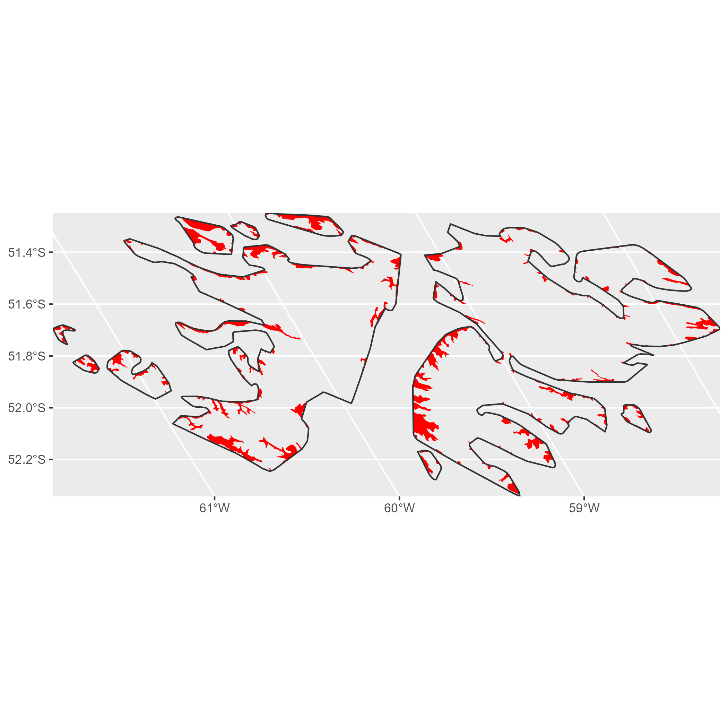

**(a)**

**(b)**

#### Human Footprint

We used the Human Footprint rasters for 1993 and 2009 from (Venter *et al*., 2016) <https://datadryad.org/stash/dataset/doi:10.5061/dryad.052q5>. We aggregated these rasters to a spatial resolution of ~5km^2^ (reducing the number of pixels from 13.4 million to ~500k) to facilitate manipulation and polygonization. The Human Footprint 2009 map assigns each pixel a numeric value from 0 (no human impact) to 50 (maximum human impact); to use these data as predictors, we binned these values into ten class categories (Figs S4.5, S4.6). We used the value at every 10^th^ percentile to define the classes (Fig S4.5) but because of the high number of pixels with a score of 0, we used the 20^th^ percentile to define the lowest class (class 1), while the two highest classes (class 9 and class 10) are defined by the 90^th^ and 95^th^ percentile (Fig S4.5). We calculated the change in human footprint (delta_hfp) by subtracting the 1993 values from the 2009 values and binned these values into five categories (Figs S4.7, S4.8).

***Figure S4.5.*** *Density of Human Footprint (2009) values from Venter et al. (2016), with the bin thresholds used in this study shown using colour (corresponding to the colours used in Fig S4.6).*

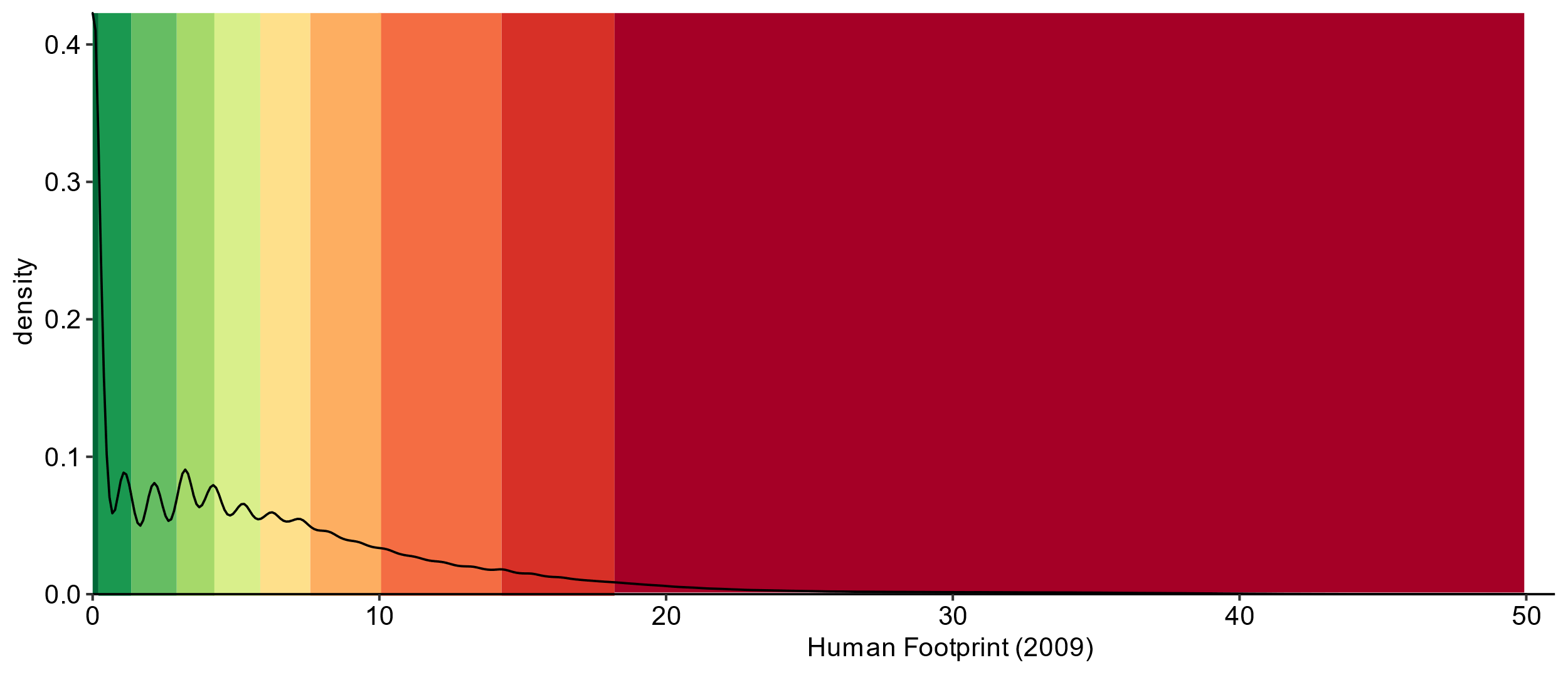

For both Human Footprint 2009 and Change in Human Footprint 1993-2009, we aggregated the raster using the classes defined above, converted it to polygons using the ‘sf’ package (Pebesma, 2018) and calculated the area of each class in each botanical country. We standardised these areas according to the same protocol for biomes (Fig S4.3b).

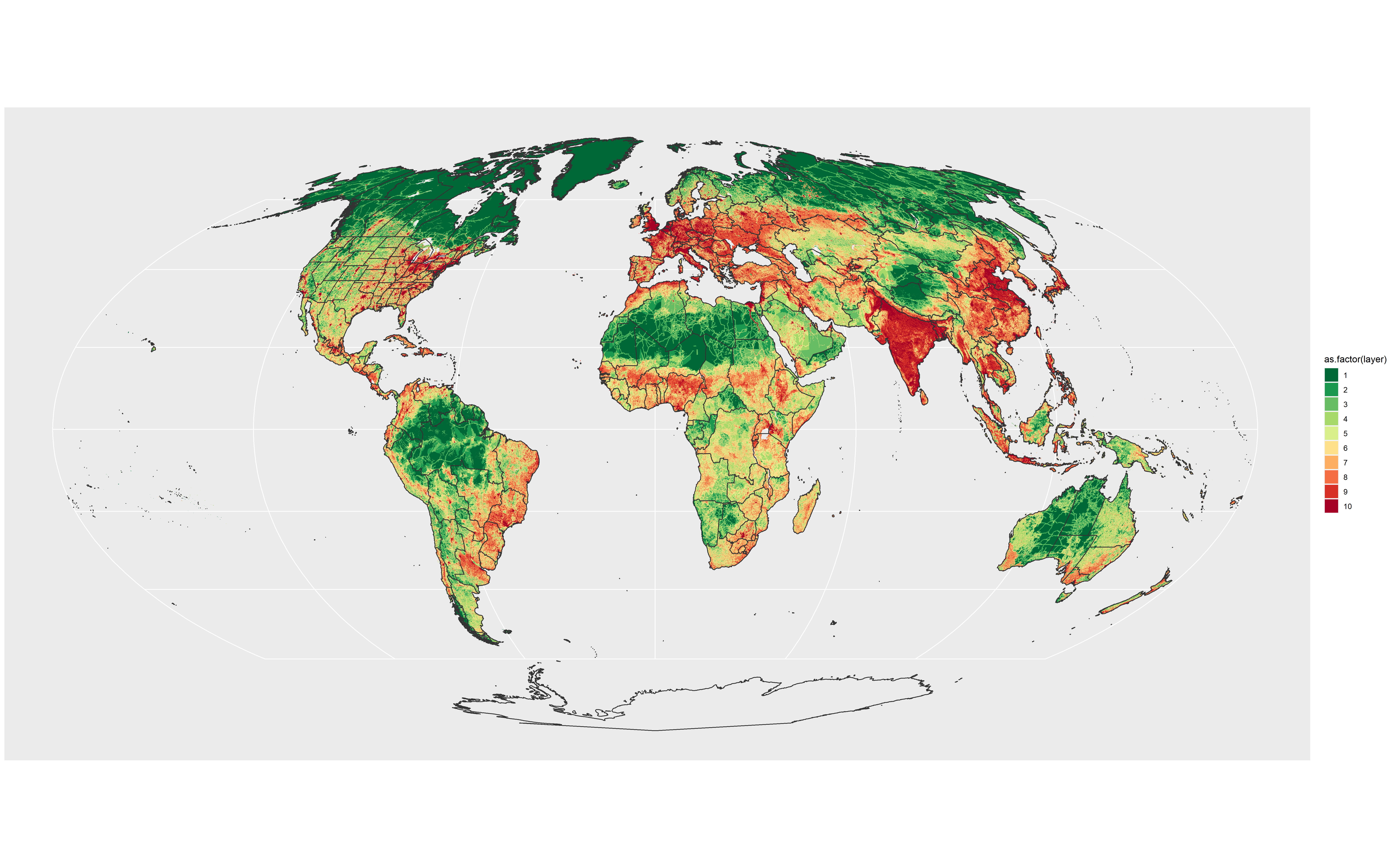

***Figure S4.6.*** *Distributions of Human Footprint (2009) classes; values binned from Venter et al. (2016) as shown in Fig S4.5*

***Figure B.*** *Density of Change in Human Footprint (1993-2009) values from Venter et al. (2016), with the bin thresholds used in this study shown using colour (corresponding to the colours used in Fig C).*

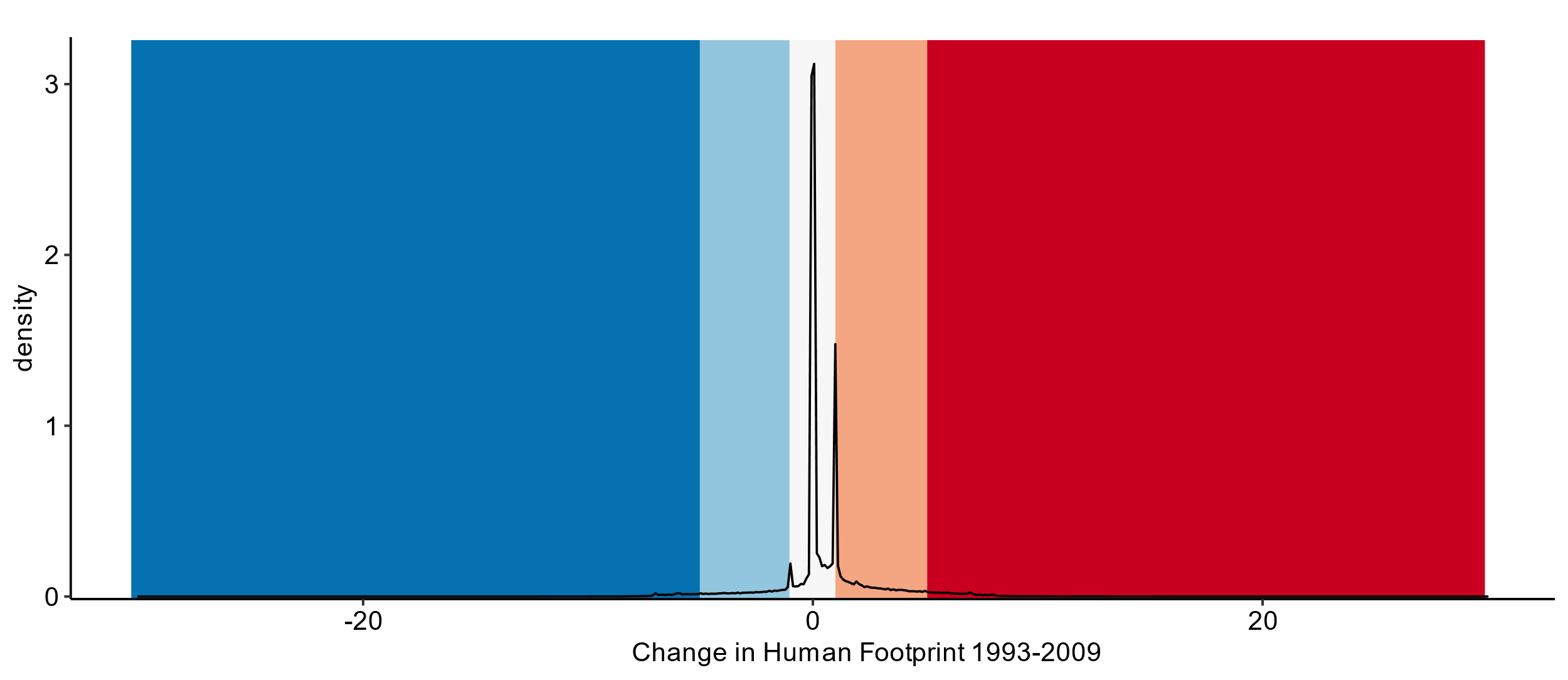

***Figure S4.7.*** *Density of Change in Human Footprint (1993-2009) values from Venter et al. (2016), with the bin thresholds used in this study shown using colour (corresponding to the colours used in Fig S4.8).*

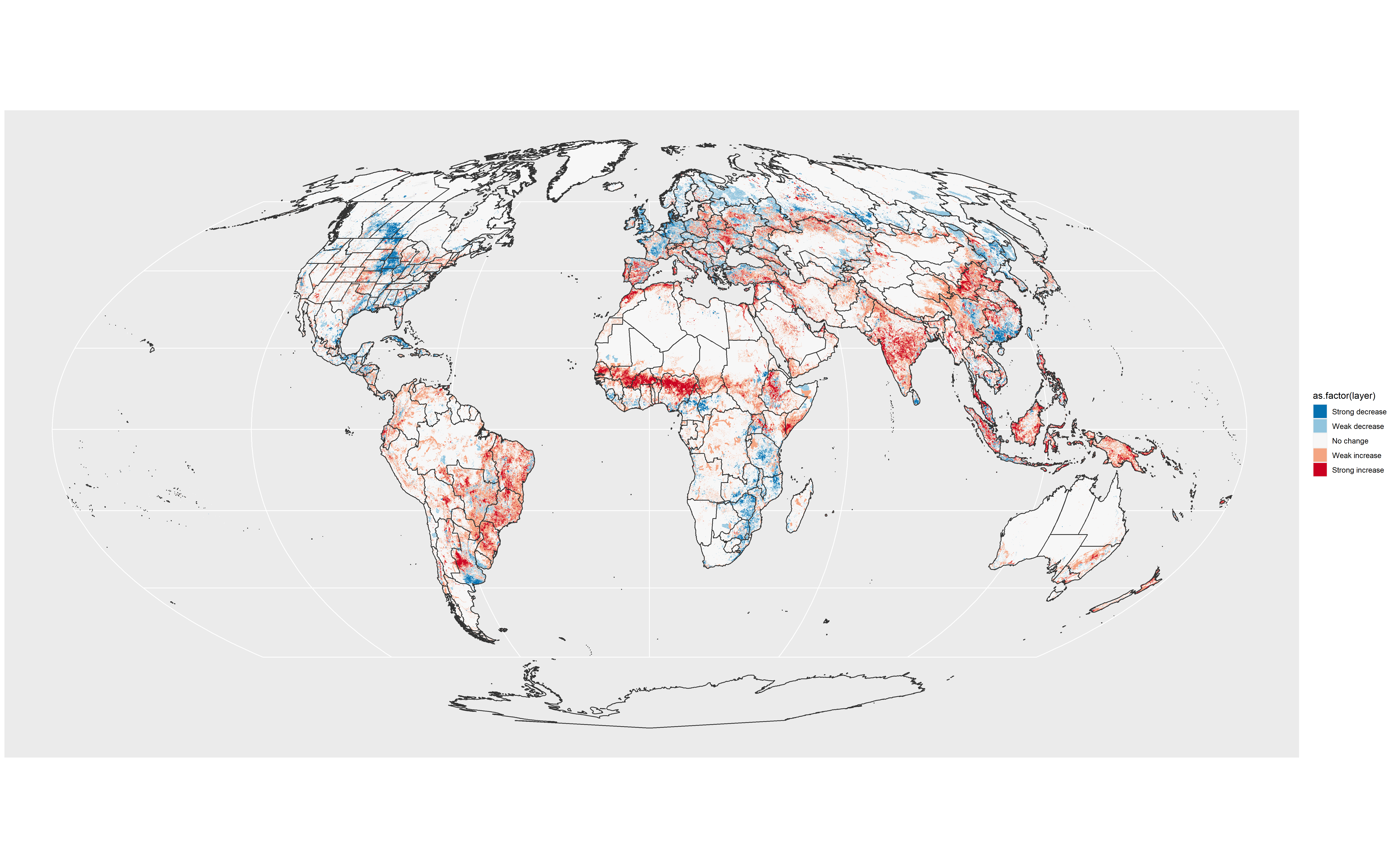

***Figure S4.8.*** *Distributions of Change in Human Footprint 1993-2009 classes; values binned from Venter et al. (2016) as shown in Fig S4.7*

There was <20% Human Footprint coverage for 46 botanical countries: small islands that range from the virtually uninhabited (e.g. Bouvet Island) to the heavily developed (Maldives). We estimated approximate percentages of Human Footprints using satellite imagery, using regions with Human Footprint coverage as references. These estimations (including justifications for assignments) are available as “Table_S6_manually_estimated_hfp2009.csv”.

### Table S5 – life_form_mapping.csv

Mapping lifeform classes used in WCVP to a simplified version with fewer classes – as used in (Humphreys et al., 2019) and modified in(Nic Lughadha et al., 2020): **“life_form_mapping.csv”**

### Table S6 – manual human footprint estimations for Human Footprint 2009

See item “Table_S6_manually_estimated_hfp2009.csv” and item S4 for explanation.

### Table S7 – comparison of model performance with other studies

| **Short name** | **Accuracy** | **Sensitivity** | **Specificity** | **TSS** | **Reference** |
| --- | --- | --- | --- | --- | --- |
| Walker_evidence_min | 0.5 |  |  |  | (Walker et al., 2022) |
| Silvestro_madagascar | 0.67 |  |  |  |  |
| This study | 0.79 | 0.76 | 0.8 | 0.58 |  |
| Walker_evidence_max | 0.8 |  |  |  | (Walker et al., 2022) |
| Bellot_palms | 0.81 | 0.77 | 0.86 | 0.63 | (Bellot et al., 2022) |
| Silva_trees | 0.83 | 0.78 | 0.87 | 0.63 | (Silva et al., 2022) |
| Zizka_orchids | 0.84 |  |  |  | (Zizka et al., 2021) |
| NicLughadha_misuse | 0.9 | 0.85 | 0.91 | 0.76 | (Nic Lughadha et al., 2019) |
| Darrah_monocots | 0.91 | 0.88 | 0.93 | 0.81 | (Darrah et al., 2017) |

### Table S8 –model performance

Based on recent Red List assessments i.e. filtered to only include the last 10 years (51,176 species) and based on unseen published assessments from Red List version 2022-2, from a model developed using the 2022-1 Red List dataset.

| **Method** | **Accuracy** | **Sensitivity** | **Specificity** | **True Skill Statistic (TSS)** |
| --- | --- | --- | --- | --- |
| Recent (<10 years) | 0.790 | 0.762 | 0.811 | 0.574 |
|  | [0.785, 0.795] | [0.757, 0.772] | [0.796, 0.820] | [0.564, 0.581] |
| Unseen assessments (2022-2) | 0.766 | 0.793 | 0.724 | 0.516 |

### Figure S9 – performance plots

Performance of the mode on different facets of the data: (A) life form, (B) Biome, (C) large families (i.e. with at least 3,000 species). Two cross-validation (CV) methods compared: random and family-wise blocks (See methods). Numbers indicate number of species in each group.

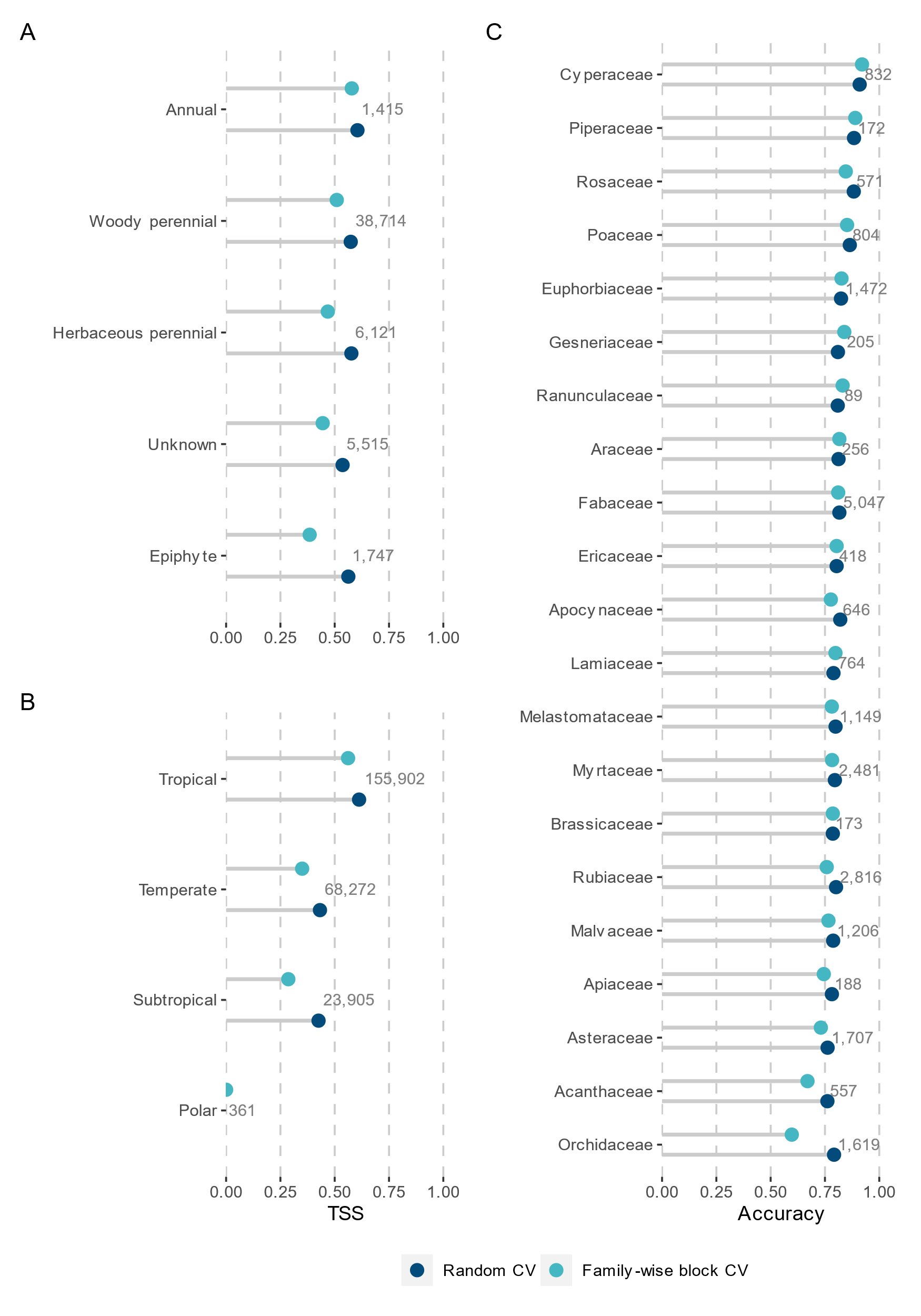

### Figure S10 – performance maps

(A) True skill statistic (TSS) scores mapped for botanical countries (mean of TSS for all species occurring in each region), (B) mean sensitivity for all species occurring in each region and (C) mean specificity for all species occurring in each region.

A high sensitivity score indicates a high proportion of the threatened species are being correctly predicted. A high specificity score indicates a high proportion of the non-threatened species are being correctly prediction. The TSS is a performance metric that balances the proportion of threatened species correctly predicted as such (sensitivity) with the proportion of non-threatened species correctly predicted (specificity). The true skill statistic (TSS) ranges from 1 for perfect predictions to -1 if there are no correct predictions. A score >= 0.5 could be considered a good model.

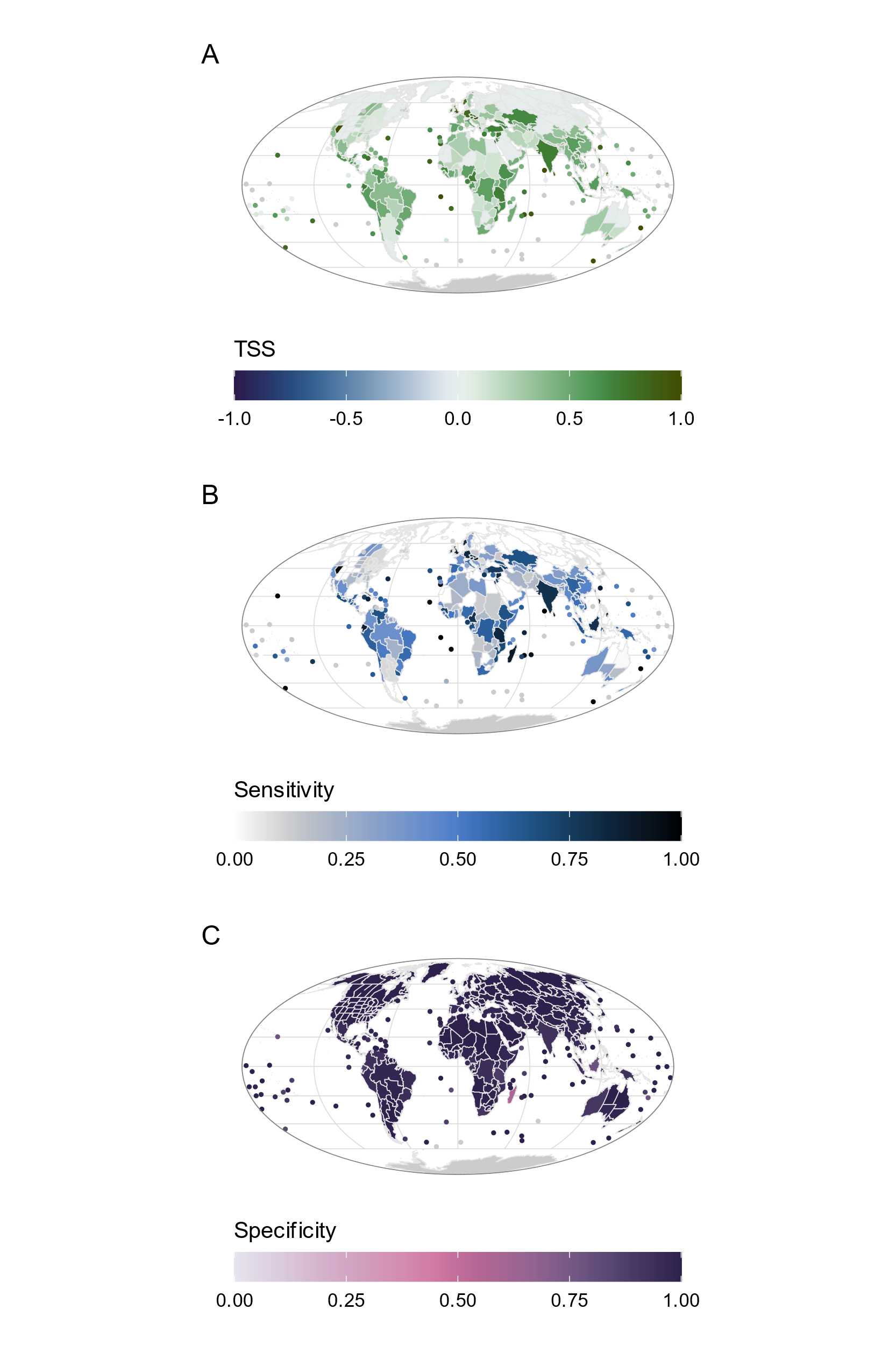

### Table S11 – performance by WGSRPD ‘continent’

|  | WGSRPD continent | | | | | | | | |
| --- | --- | --- | --- | --- | --- | --- | --- | --- | --- |
|  | Europe | Africa | Asia Temp. | Asia Trop. | Australia | Pacific | Northern America | Southern America | Antarctic |
| accuracy | 0.978 | 0.932 | 0.963 | 0.929 | 0.911 | 0.940 | 0.972 | 0.929 | 0.834 |
| TSS | 0.346 | 0.395 | 0.275 | 0.499 | 0.331 | 0.339 | 0.124 | 0.399 | 0.542 |
| sensitivity | 0.353 | 0.429 | 0.288 | 0.536 | 0.367 | 0.400 | 0.129 | 0.424 | 0.463 |
| specificity | 0.993 | 0.966 | 0.987 | 0.965 | 0.967 | 0.963 | 0.995 | 0.975 | 0.972 |

Temperate Asia and Northern America had high specificity scores (highlighted in green) but were the regions with the lowest TSS scores due to the low sensitivity (highlighted in blue).

### Figure S12 – median range size vs sensitivity

Sensitivity scores of species against median range size of species (number of botanical countries).

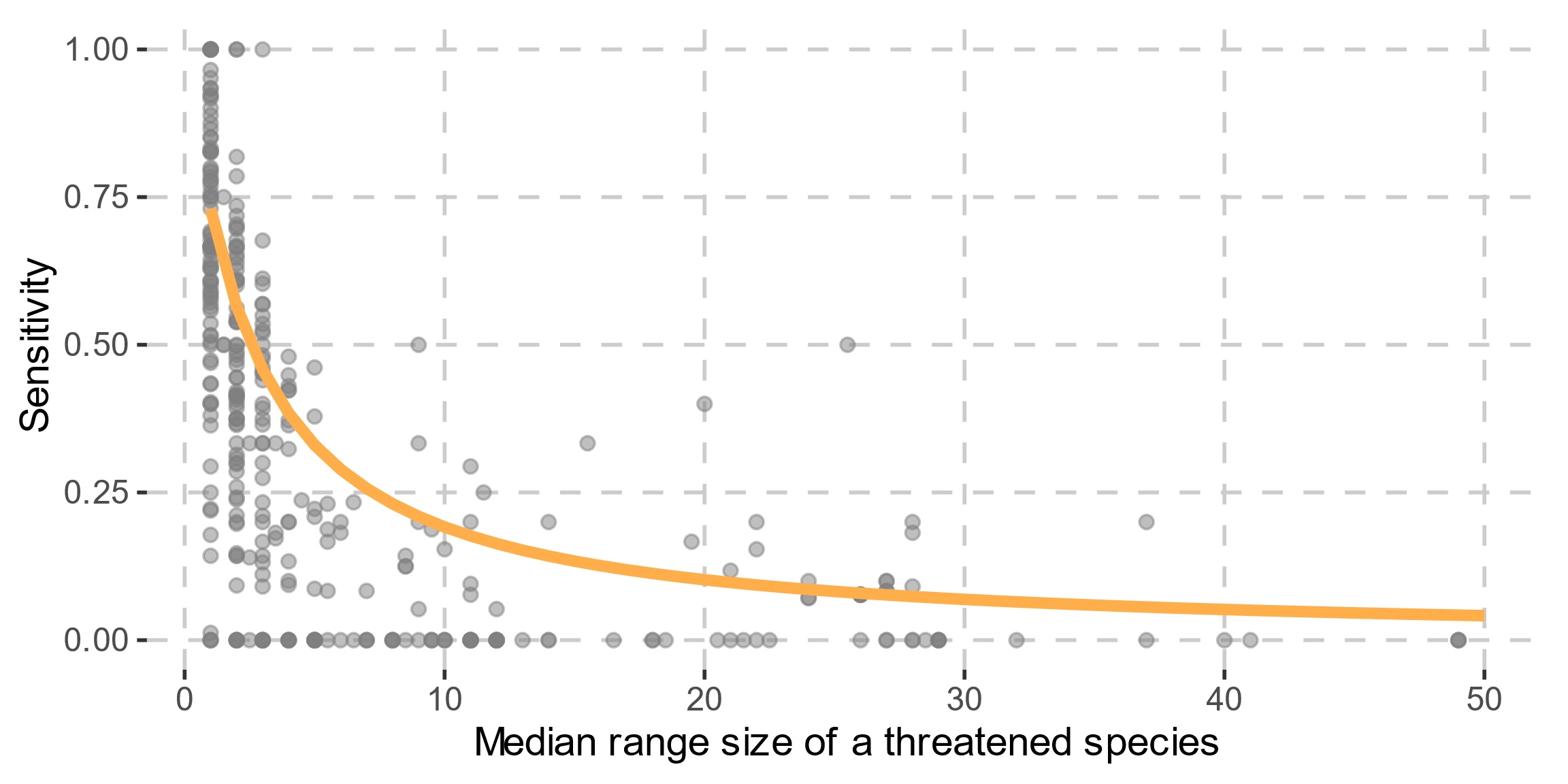

### Figure S13 – sensitivity vs citation of different Red List criteria

Breakdown of Red List criteria cited for threatened species (%) and sensitivity scores for threatened species cited or not cited per criterion (A, B, C and D).

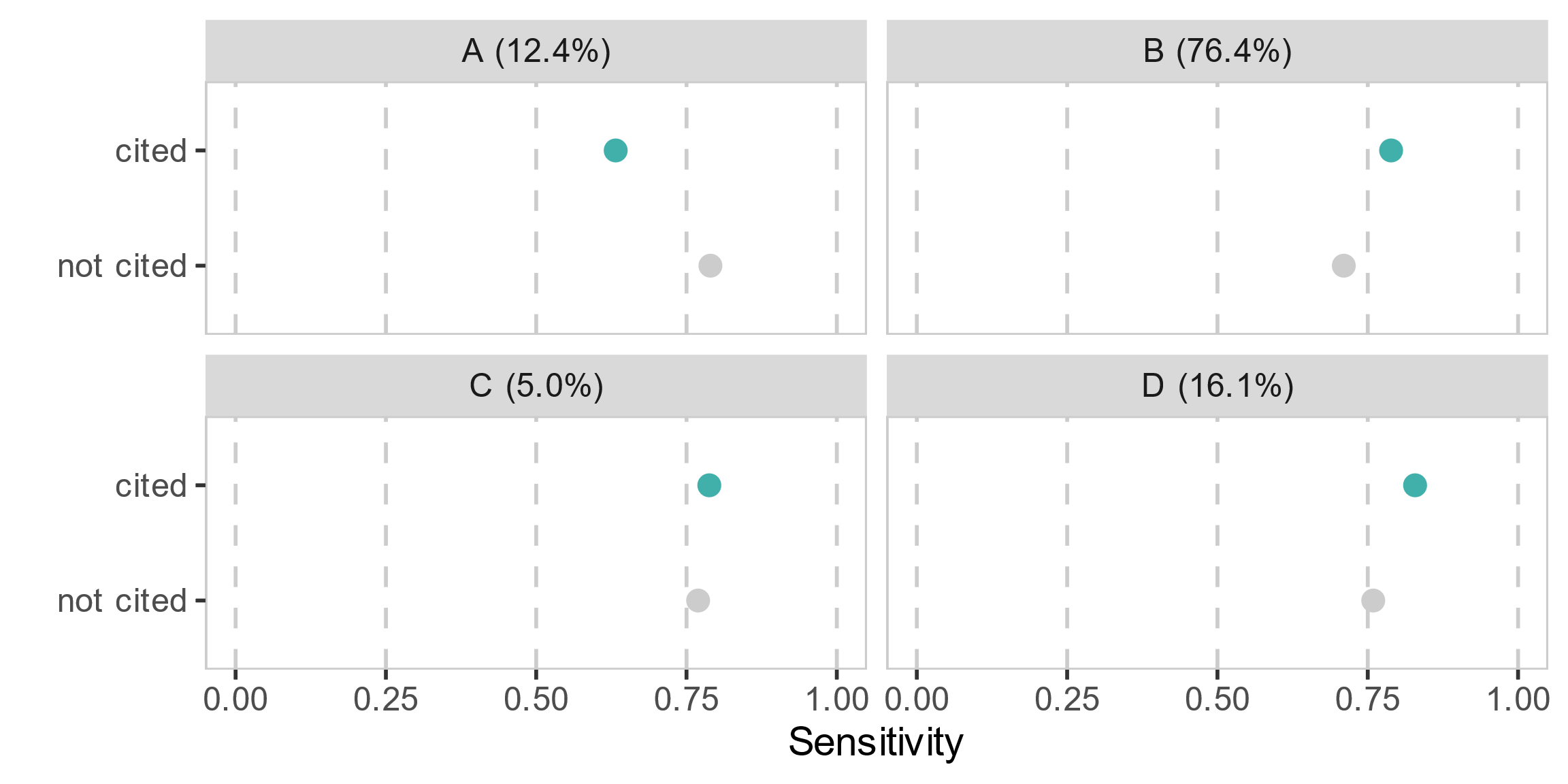

### Figure S14 – sensitivity vs citation of criterion B1, B2 or both

Average sensitivity of threatened species that cited sub-criterion B1, B2, or both, with confidence intervals.

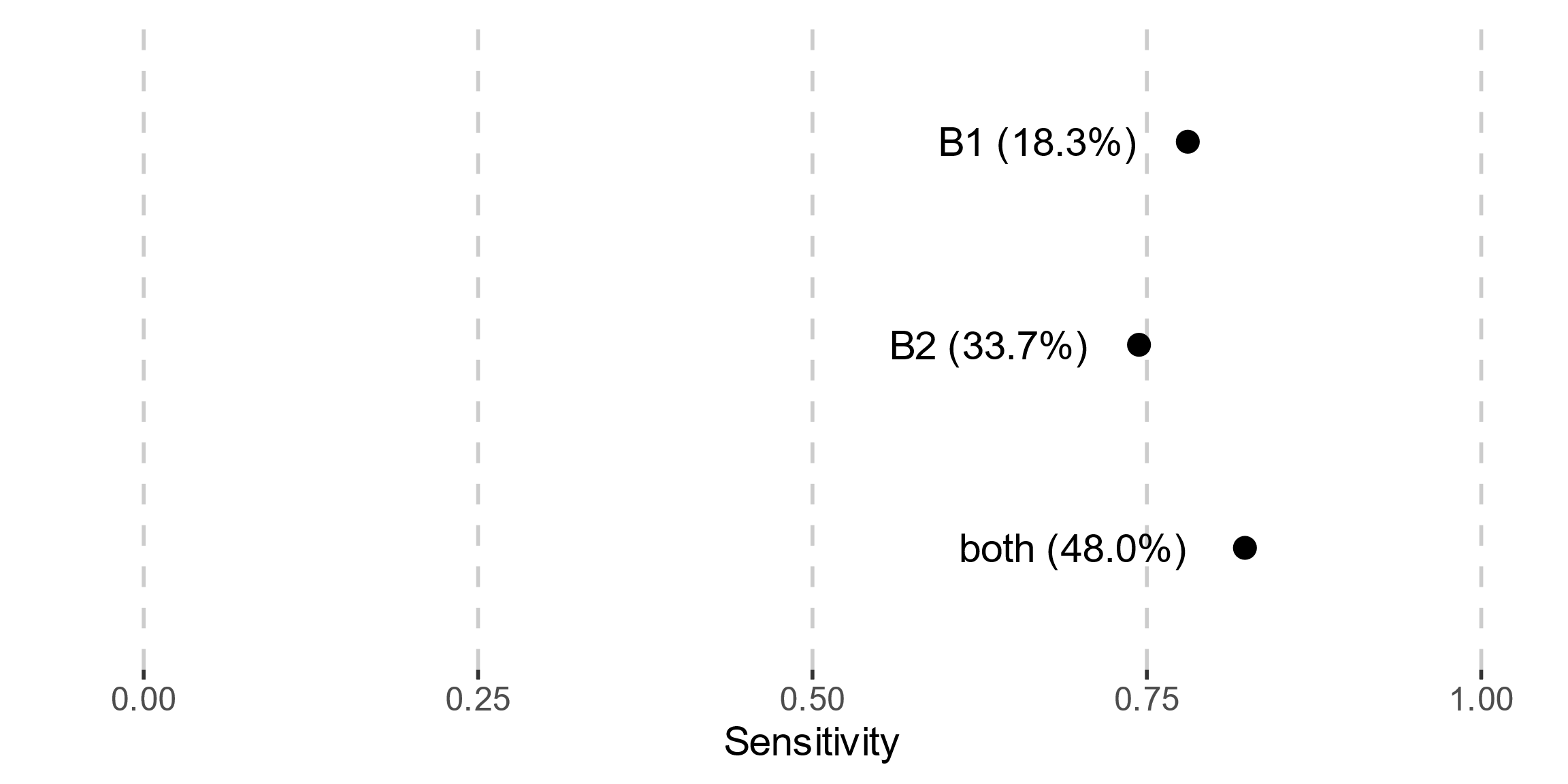

### Figure S15 – threats

Summary of threats coded using IUCN threats classification. Species that were correctly classified (correct) compared against those that were not correctly classified (incorrect).

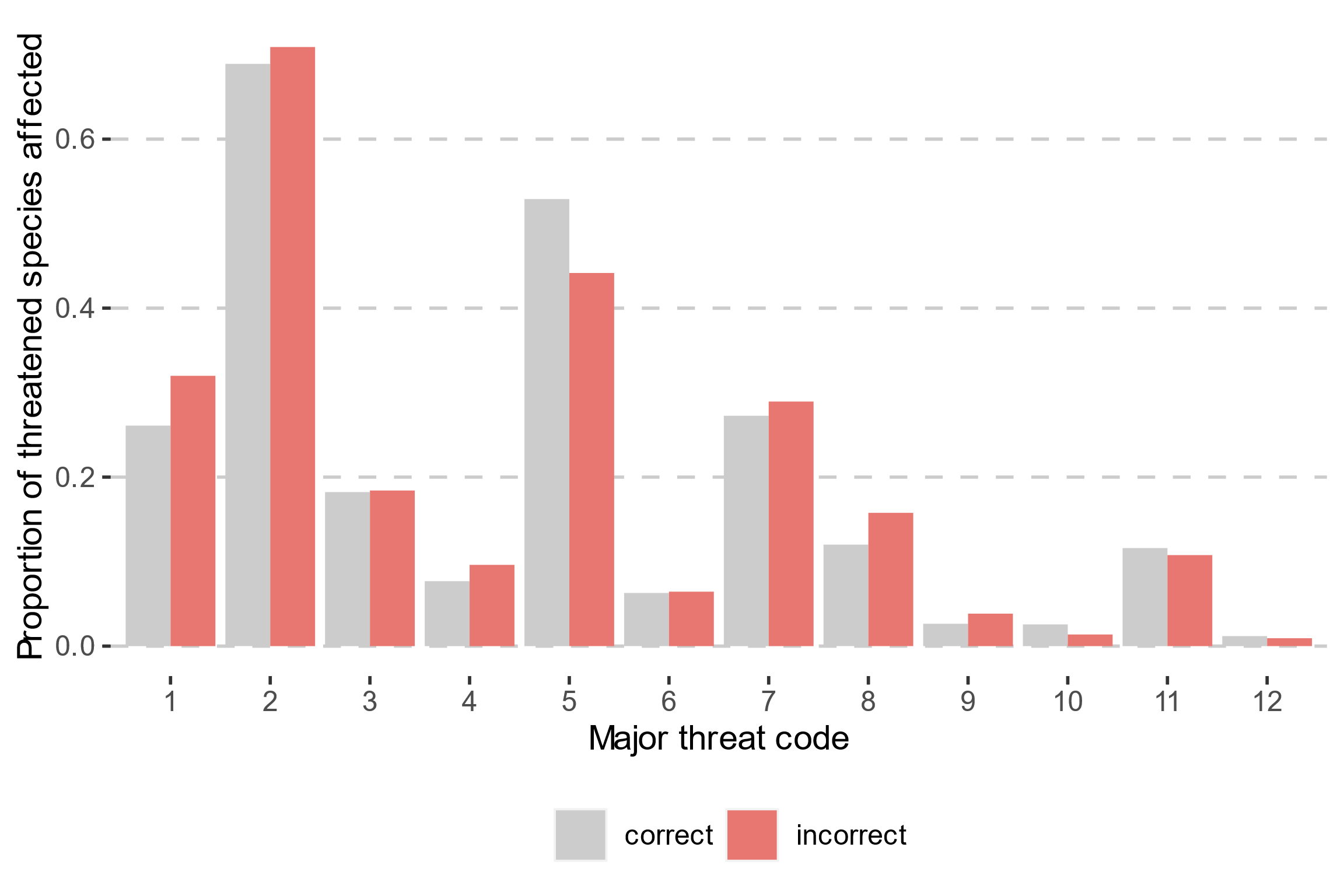

Threat codes conversion:

1 = Residential & commercial development

2 = Agriculture & aquaculture

3 = Energy production & mining

4 = Transportation & service corridors

5 = Biological resource use

6 = Human intrusions & disturbance

7 = Natural system modification

8 = Invasive & other problematic species

9 = Pollution

10 = Geological events

11 = Climate change & severe weather

12 = Other options

### Table S16 – final predictions

Species level predictions with predicted class and uncertainty values: **“bart-final-predictions.csv”**

### Figure S17 – predictor importance plot

Importance of individual predictors measured by mean decrease in accuracy when the values of each predictor are randomly shuffled. For descriptions of predictors see Table S3.

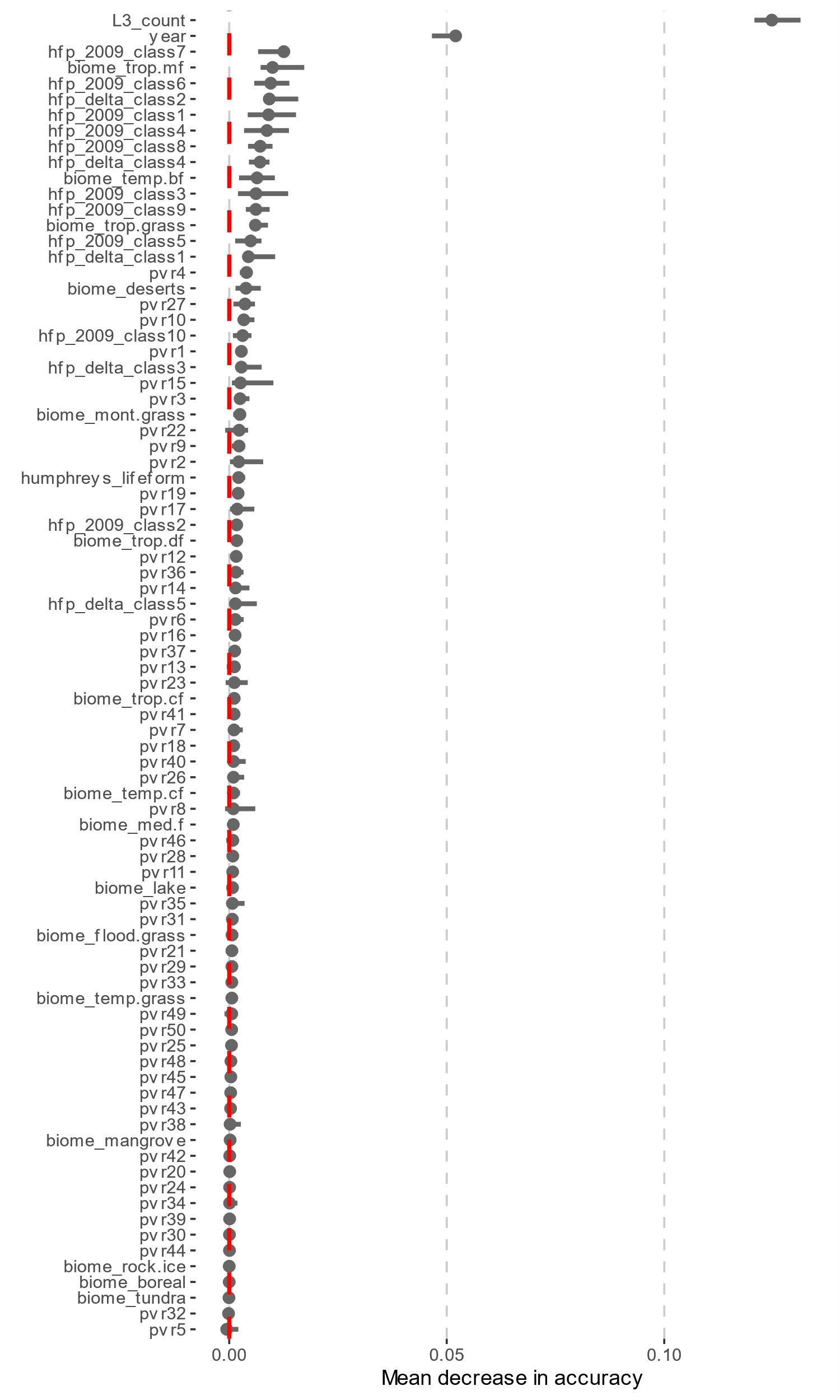

### Figure S18 – red list summary statistics

Summary statistics for angiosperms on the Red List version 2022.2.

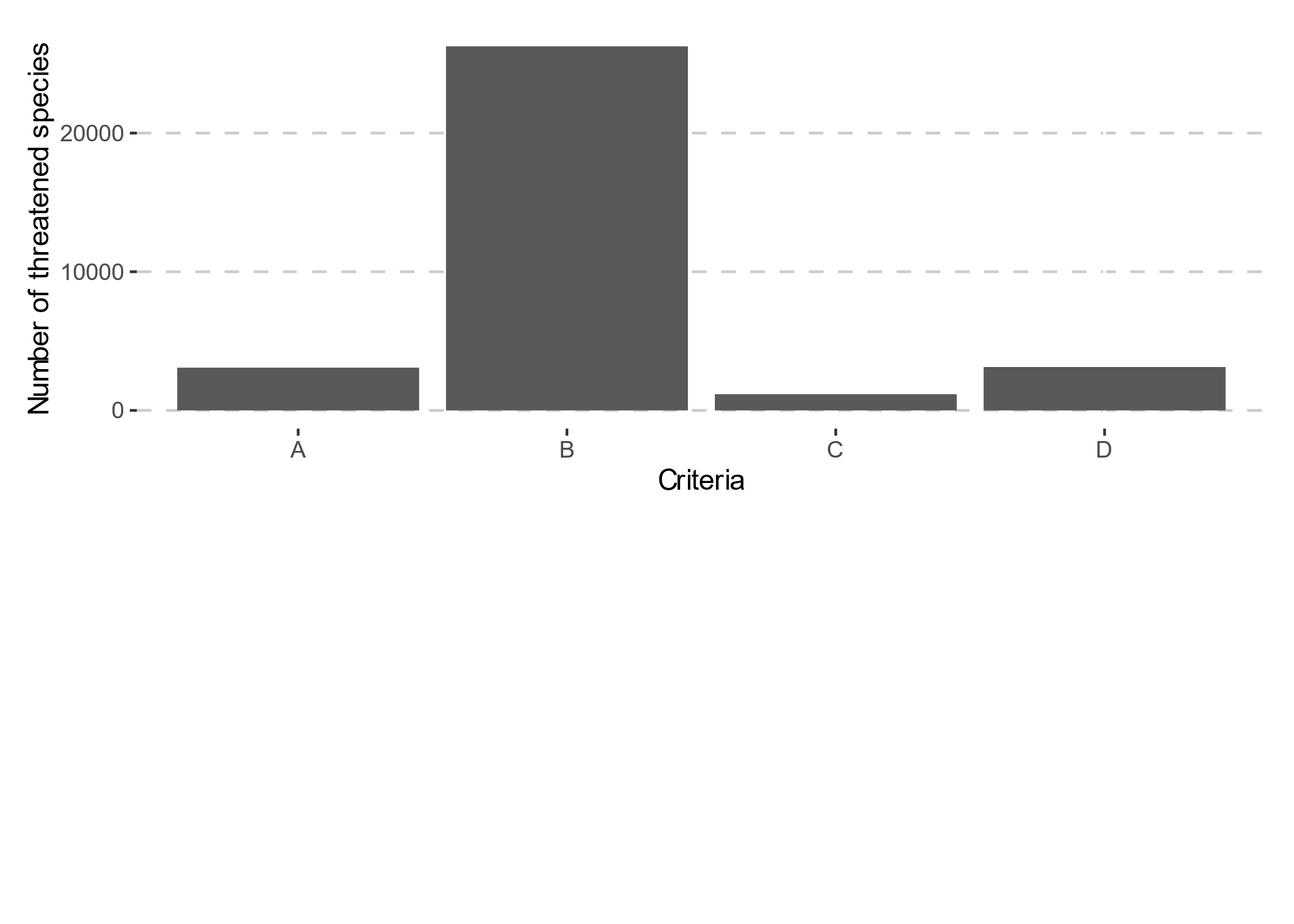

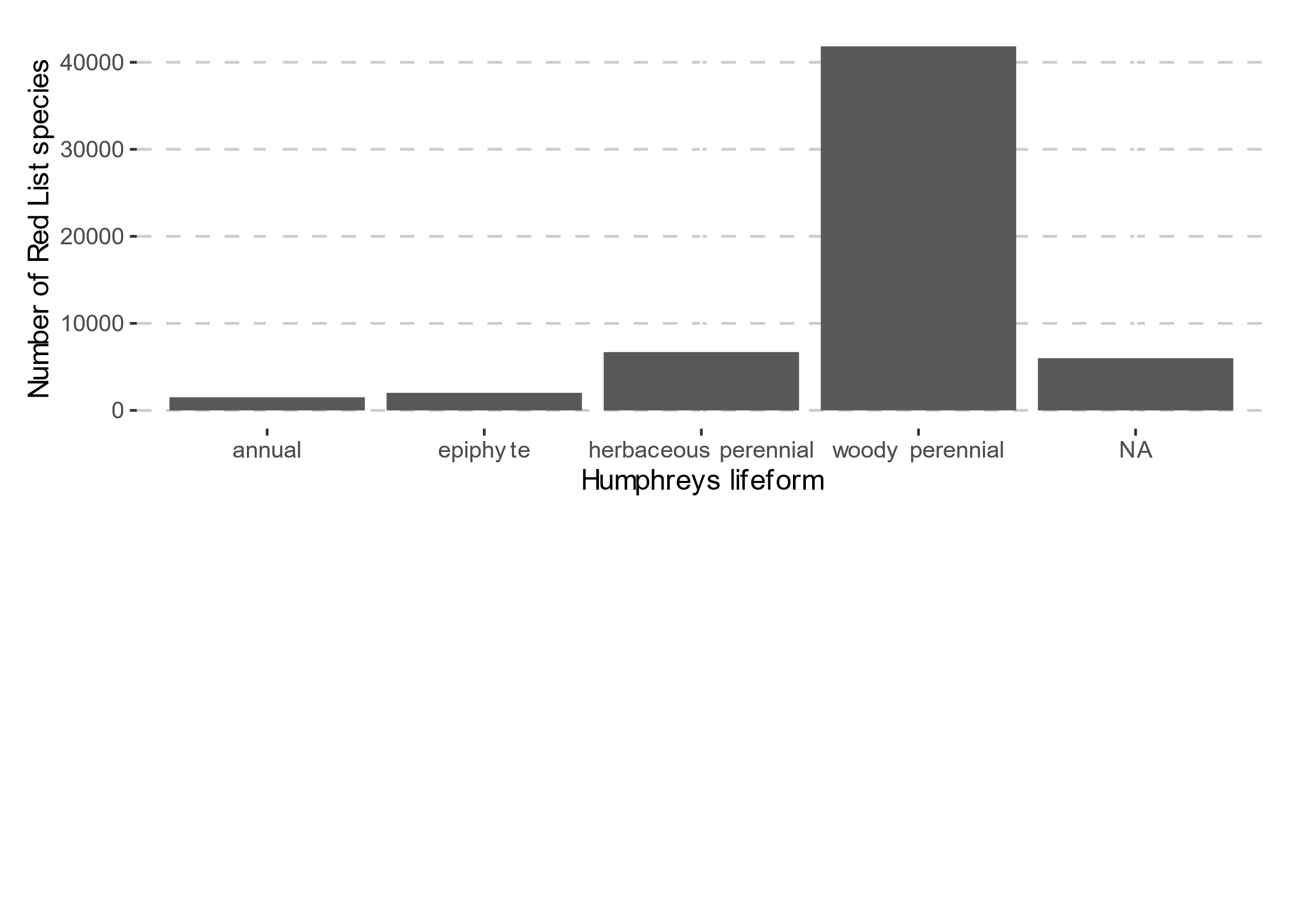

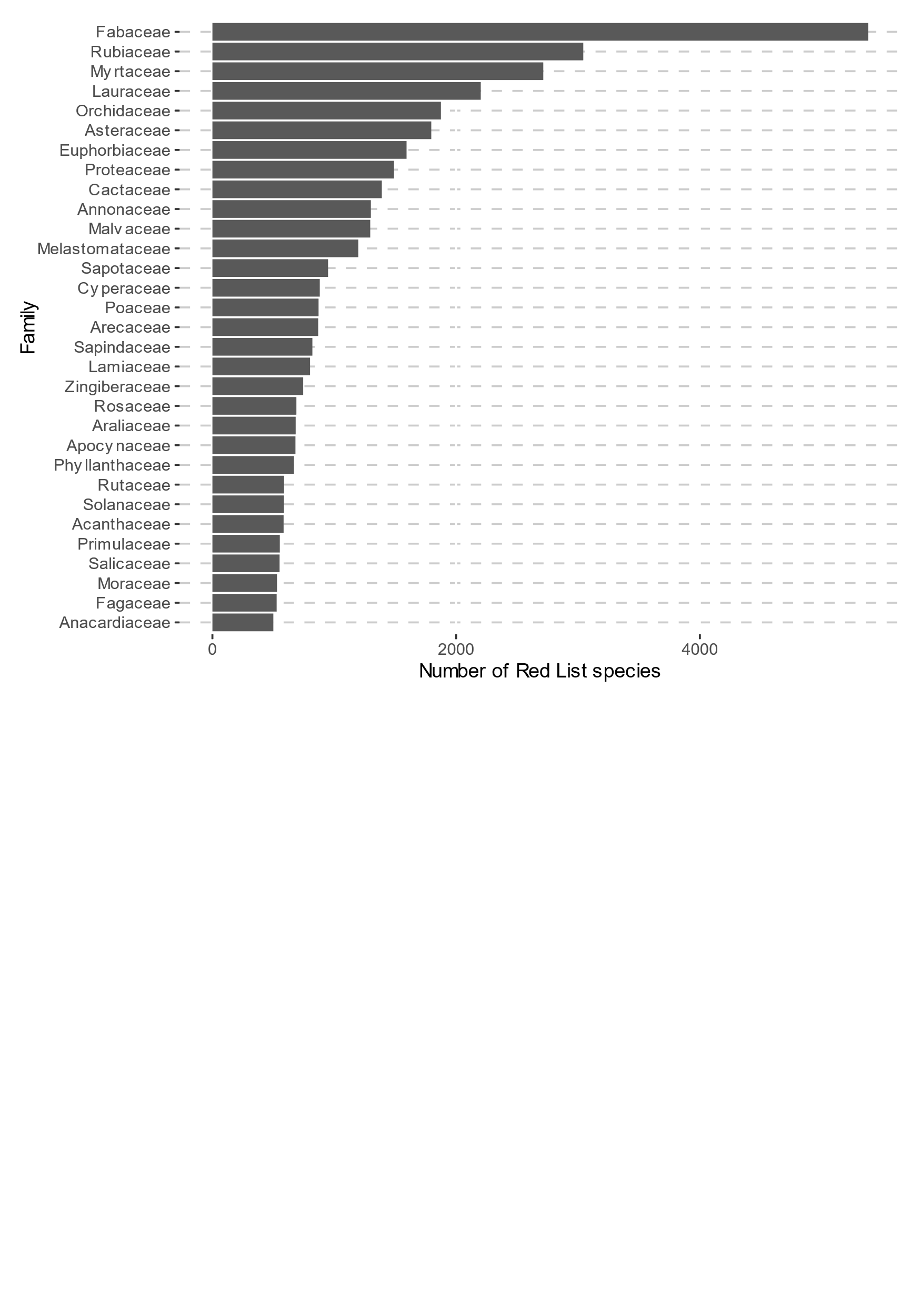

### Figure S19 –Red List criteria over time

Number of plant assessments categorised as threatened or Near Threatened using different Red List criteria (A, B, C and D) over time.

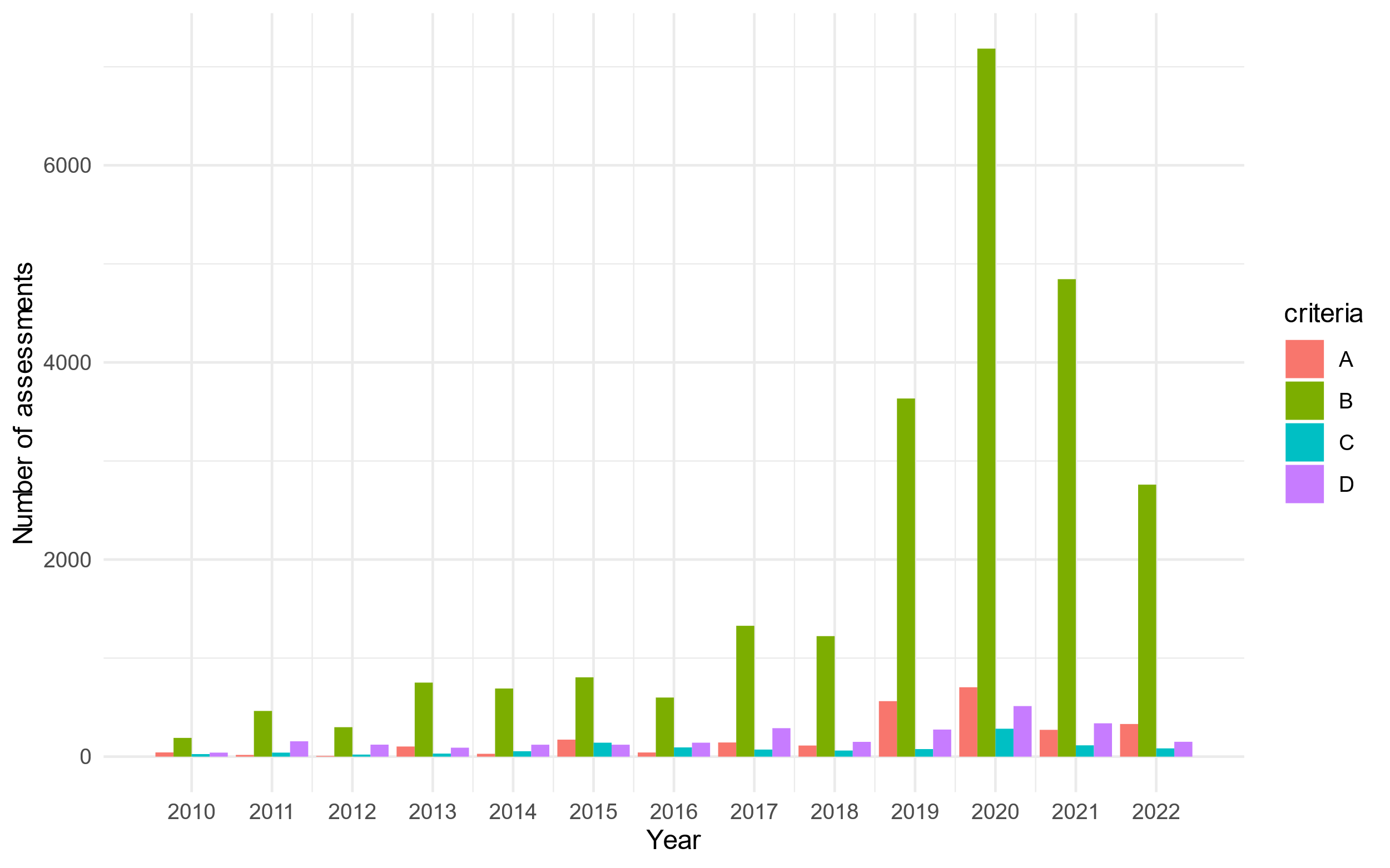

### Figure S20 – criteria combinations

Threatened angiosperm species assessments coded with different combinations of Red List criteria (combinations of A, B, C and D) – count of assessments to the right of columns. Mean threat level was calculated as mean threat score for all assessments under each criterion combination where threat categories were converted to a number CR = 5, EN = 4, VU = 3, NT = 4.

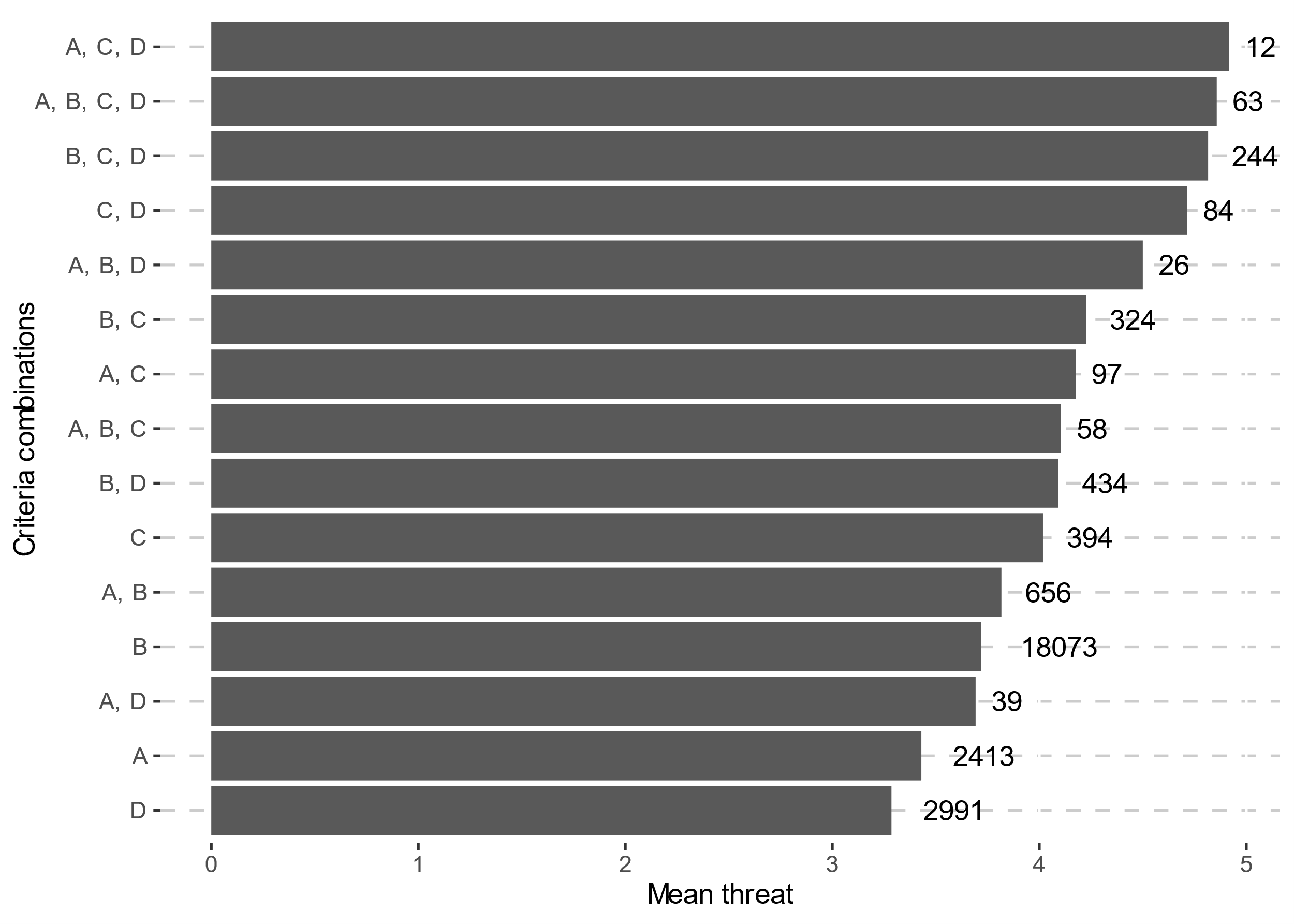

### Table S21 – Summary of performance, importance, and predictions.

See table **S21_** **bart-summary.xlsx** for summary tables of performance, importance and predictions.
